# IGSF3 is a homophilic cell adhesion molecule that drives lung metastasis of melanoma by promoting adhesion to vascular endothelium

**DOI:** 10.1101/2023.10.16.562470

**Authors:** Yue Guo, Yutaka Kasai, Yuto Tanaka, Yuki Ohashi-Kumagai, Takeshi Ito, Yoshinori Murakami

## Abstract

The immunoglobulin superfamily (IgSF) is one of the largest families of cell-surface molecules involved in various cell-cell interactions, including cancer–stromal interactions. In this study, we conducted a comprehensive RT-PCR-based screening for IgSF molecules that promote experimental lung metastasis in mice. By comparing the expression of 325 genes encoding cell-surface IgSF molecules between mouse melanoma B16 cells and its highly metastatic subline, B16F10 cells, we found that expression of the *Immunoglobulin superfamily member 3* (*Igsf3*) was significantly enhanced in B16F10 cells than in B16 cells. Knockdown of *Igsf3* in B16F10 cells significantly reduced lung metastasis following intravenous injection into C57BL/6 mice. IGSF3 promoted adhesion of B16F10 cells to vascular endothelial cells and functioned as a homophilic cell adhesion molecule between B16F10 cells and vascular endothelial cells. Notably, the knockdown of IGSF3 in either B16F10 cells or vascular endothelial cells suppressed the transendothelial migration of B16F10 cells. Moreover, IGSF3 knockdown suppressed the extravasation of B16F10 cells into the lungs after intravenous injection. These results suggest that IGSF3 promotes the metastatic potential of B16F10 cells in the lungs by facilitating their adhesion to vascular endothelial cells.

## Introduction

Metastasis, the primary cause of cancer-related deaths in solid tumors, is a multi-step process that includes invasion of the blood or lymphatic vessels, survival within the circulation, extravasation to distant organs, and subsequent proliferation at these sites ^1^. Metastasis is contributed to not only by tumor cells but also by tumor-associated stromal cells, which constitute the tumor microenvironment ^2^. Interactions between tumor and host cells are pivotal at each step of metastasis. For instance, cancer cells can evade immune cell attacks within the circulation and attach to vascular endothelial cells (vECs) in the first step of extravasation. The importance of tumor-host interactions in metastasis has been extensively studied using mouse models, notably the model of experimental lung metastasis with mouse melanoma B16F10 cells. A variety of host factors, including immune regulators and cell-surface molecules, are essential in metastasis ^3, 4^. However, a comprehensive understanding of the mechanisms underlying this complex process is lacking.

The immunoglobulin superfamily (IgSF) is the largest family of cell-surface and secreted molecules that mediate cell-cell interactions in immune regulation, cell adhesion, and neural transmission. IgSF is characterized by molecules containing immunoglobulin (Ig)-like loops and consists of approximately 500 members in humans, excluding T cell receptors and immunoglobulins that undergo gene rearrangement ^5^. Members of the IgSF range from cell adhesion molecules to immune co-inhibitory/stimulatory molecules, and receptor tyrosine kinases/phosphatases. They exert diverse functions through homophilic interactions or heterophilic interactions with other IgSF molecules, integrins, and proteins with different domain types ^6^. IgSF-mediated interactions, represented by the CTLA-4–B7 interaction in immune checkpoint regulation ^7^ and the VCAM1–integrin interaction in cancer cell adhesion to vECs ^8^, have been linked to metastasis. However, despite the evident roles of IgSF interactions in metastasis, a systematic screening of IgSF molecules that regulate metastasis has not yet been performed.

In this study, we conducted RT-PCR-based screening to identify IgSF molecules implicated in the experimental lung metastasis of B16F10 cells. Candidate metastasis-promoting genes were selected based on their elevated expression in highly metastatic B16F10 cells compared to that in their parental B16 counterparts. On the basis of the results of experimental lung metastasis with B16F10 cells after the knockdown of each gene, we identified IGSF3 as a metastasis promoter that mediates tumor-endothelium interactions and as a potential prognostic factor for human melanoma.

## Materials and Methods

### Cell lines and antibodies

The cell lines and antibodies used are described in Document S1.

### RT-PCR

Total RNA was extracted from cells using an RNeasy Mini Kit (QIAGEN, Venlo, Netherlands) and reverse-transcribed using a ReverTra Ace qPCR RT Kit (TOYOBO, Osaka, Japan). RT-PCR was performed using KOD FX DNA polymerase (TOYOBO). Quantitative RT-PCR was performed using CFX Connect Real-Time PCR System (Bio-Rad, Hercules, CA, USA) with PowerUp SYBR Green Master Mix (Thermo Fisher Scientific, Waltham, MA, USA). The primer sets for the genes encoding mouse IgSF are listed in Table S1. Mouse *Gapdh* served as an internal control: sense, 5’-CCACTCTTCCACCTTCGATG-3’; antisense, 5’-GGAGGGAGATGCTCAGTGTTG-3’. The relative gene expression was determined using the 2^-ΔΔCT^ method.

### Lentiviral vectors

For knockdown, Mission shRNA clones were purchased from Sigma-Aldrich. The sequences of each shRNA are presented in Table S2. The control vectors, pLKO.1 and pLKO.5 were sourced from Addgene (#1864; a gift from David Sabatini ^9^) and Sigma-Aldrich (St. Louis, MO, USA), respectively. For overexpression, IGSF3 and EGFP expression vectors were generated using the pLenti6-V5/DEST vector (Thermo Fisher Scientific). The coding region of human *IGSF3* was amplified using the primer sets listed in Table S3, cloned into the pENTR/D-TOPO vector (Thermo Fisher Scientific), and subsequently recombined with pLenti6-V5/DEST using LR clonase (Thermo Fisher Scientific). The EGFP sequence was integrated into pLenti6-V5/DEST at the *SpeI*-*SacII* site using Gibson Assembly (New England Biolabs, Ipswich, MA, USA). These lentiviral vectors were infected to cell lines as previously described ^10^. Stable cell lines were selected using either puromycin (1 μg/mL) or blasticidin (10 μg/mL). EGFP-expressing cells were further sorted using FACSAria III (BD Biosciences, Franklin Lakes, NJ, USA).

### RNA interference

MS1 cells were seeded in 24-well plates and cultured until they reached 50% confluency. The cells were then transfected with either the siRNA SMARTpool for mouse *Igsf3* (Horizon Discovery, Cambridge, UK) or the ON-TARGET plus siControl Non-targeting Pool (Horizon Discovery) using the Lipofectamine RNAiMAX Transfection Reagent (Thermo Fisher Scientific).

### Animal experiments

Female C57BL/6J mice (CLEA Japan, Tokyo, Japan) aged 6–8 weeks were used for the experiments. The experimental lung metastasis assay involved injecting a suspension of 5 × 10^4^ cells in 200 μL PBS into the tail vein. The mice were euthanized two weeks after injection, and the number of metastatic foci formed on the lung surface was counted. For extravasation studies, mice were injected with 2 × 10^5^ EGFP-expressing cells in 200 μL of PBS into the tail vein. After 24 h, the extracted lungs were immunostained with anti-CD31 antibody to visualize vascular endothelial cells as previously described ^11^. The subcutaneous tumor formation assay was performed by injecting a suspension of 5 × 10^5^ cells in 200 μL PBS into both flanks of the mice. Tumor diameters were measured twice a week, and the tumor volume was calculated as 1/2 × length × width^2^. All animal experiments were performed in accordance with the guidelines of The Institute of Medical Science, The University of Tokyo.

### Cell growth assay

The cells were seeded in 6-well plates at a density of 5 × 10^3^ cells/well. The number of cells cultured for two to five days was counted using a hemocytometer (Funakoshi, Tokyo, Japan).

### Soft agar colony formation assay

A suspension of 2 × 10^3^ cells in a growth medium containing 0.33% Noble agar (BD Biosciences) was plated onto 6-well plates precast with 0.5% Noble agar. After two weeks of incubation, the colonies were counted using a stereomicroscope (Olympus, Tokyo, Japan).

### Endothelium adhesion assay

MS1 cells were seeded in 24-well plates at a density of 8 × 10^4^ cells/well and cultured for 24 h to form a monolayer. Subsequently, B16F10 cells were labeled with Calcein-AM (Fujifilm Wako Pure Chemical, Osaka, Japan) at 2 µM for 30 min, and 8 × 10^3^ cells were overlaid onto the MS1 monolayer. After incubation for five minutes, the unattached cells were washed off with PBS. The number of B16F10 cells that adhered to the MS1 cells was counted after fixation with 4% paraformaldehyde (PFA) in ten randomly selected microscopic fields using a fluorescence microscope (BZ-X700; Keyence, Osaka, Japan).

### Transwell migration assay

For migration assay, 5 × 10^4^ cells in serum-free medium were seeded into Transwell inserts (8.0-μm pore, Corning, NY, USA), with growth medium containing 10% FBS added to the lower chamber. After 24 h of incubation, the migrated cells were fixed with 4% PFA, stained with 0.1% crystal violet, and imaged using a microscope (BZ-X700). For transendothelial migration assay, MS1 cells were seeded in the Corning BioCoat Matrigel Invasion Chamber (8.0-µm pore, Corning) at a density of 5 × 10^4^ cells/well and cultured for 24 h to form a monolayer. Then, 1 × 10^5^ calcein-AM-labeled B16F10 cells suspended in serum-free medium were overlaid onto the MS1 monolayer, and a growth medium containing 10% FBS was added to the lower chamber. After an 8 h of incubation, the migrated cells were fixed and imaged using a microscope. The number of migrating cells waw counted using the ImageJ software (NIH, Bethesda, MD, USA).

### Wound healing assay

A suspension of 1.4 × 10^4^ cells was inserted into each well of Culture-Insert in µ-Dish (Ibidi, Martinsried, Germany) and incubated for 24 h. After removing the insert and adding the culture medium, the wound areas were phographed at 0, 12, and 24 h and measured using the ImageJ software.

### Production of IGSF3 proteins

The extracellular domain (ECD) of IGSF3 and its deletion mutants, Ig1–Ig4, Ig5–Ig8, Ig1–Ig5, Ig1–Ig6, and Ig1–Ig7 were amplified using the primer sets listed in Table S3 and cloned into pENTR/D-TOPO. The production of IGSF3 ECD with a Cd4-FLAG-His tag or a human IgG_2_-Fc tag is described in Document S1.

### Flow cytometry-based binding assay

IGSF3-Fc or normal human IgG (250 ng) and APC-conjugated anti-human IgG-Fc antibody (BioLegend, San Diego, CA, USA) was added to 1 × 10^5^ of MS1 cells suspended in 100 µL of FACS buffer (0.5% BSA/HBSS) in round bottom tubes with a volume of 5 mL (Corning). After incubation for 30 min in the dark, cells were washed with 2 mL of FACS buffer and resuspended in 500 µL of FACS buffer containing 0.1% propidium iodide. The binding of IGSF3-Fc to the cell surface was detected as the APC fluorescence intensity using FACSVerse (BD Biosciences).

### Alpha technology

IGSF3-Cd4-FLAG-His and IGSF3-Fc proteins were prepared at concentrations of 400 nM and 1.2 µM, respectively, in PBS containing 0.2% BSA (Fujifilm Wako Pure Chemical). Each protein (5 µL) was mixed and incubated for 1 h in a 384-well AlphaPlate (PerkinElmer, Waltham, MA, USA). After diluting Alpha Screen Protein A Acceptor Beads (PerkinElmer) and Anti-FLAG Alpha Donor Beads (PerkinElmer) to 80 µg/mL in 0.2% BSA/PBS, each bead (5 µL) was mixed and incubated for 1 h in the dark (total volume = 20 µL). Molecular interactions were detected as light emissions using an EnSight Multimode Plate Reader (PerkinElmer).

### Immunoprecipitation

Expression vectors with C-terminal V5 or FLAG tags were obtained by Gateway recombination of pENTR/D-TOPO with pHEK-V5 or pHEK-FLAG destination vectors ^10^. Immunoprecipitation was performed as previously described ^12^. Briefly, the expression vectors IGSF3-FLAG, IGSF3-V5 and IGSF4 (CADM1)-V5 were transfected into 293FT cells. After 24 h, the cells transfected with IGSF3-FLAG were co-cultured with cells transfected with IGSF3-V5 or IGSF4-V5 at a 1:1 ratio. After an additional 24 h, the co-cultured cells were treated with 10 mM DTSSP (Fujifilm Wako Pure Chemical). Cells were lysed with TNE buffer and the lysates were immunoprecipitated using anti-FLAG M2 affinity gel (Sigma-Aldrich). After washing the gel, immunoprecipitated complexes were eluted with 150 μg/mL of 3 × FLAG peptide (ApexBio, Houston, TX, USA) and subjected to Western blotting.

### Chemical crosslinking

The cells were trypsinized, resuspended in PBS at a density of 5 × 10^5^ cells/mL, and incubated in the presence or absence of 3 mM bis(sulfosuccinimidyl) suberate (BS^3^; Fujifilm Wako Pure Chemical) for 30 min with rotation. Crosslinking was quenched by adding 20 mM Tris-HCl (pH 7.4). The cell suspension was examined using phase-contrast microscopy before and after the chemical crosslinking reaction to ensure single-cell status. The cells were then subjected to Western blot analysis.

### Database analysis

Gene expression and prognostic analyses of human melanoma datasets were performed using the R2 Genomics Analysis and Visualization Platform (http://r2.amc.nl). To compare IGSF3 expression in normal skin and melanoma, the GSE15605 ^13^ and GSE46517 ^14^ datasets were analyzed. For prognostic analysis, the GSE19234 ^15^ and TCGA-SKCM ^16^ datasets were used.

### Statistical Analysis

Statistical analyses were conducted using the GraphPad Prism 7 software (GraphPad Software, La Jolla, CA, USA). *P*-values were estimated using a two-sided *t*-test or the Mann–Whitney *U* test. Data are presented as mean ± standard deviation (SD). *P*-values < 0.05 were considered statistically significant.

## Results

### Screening of IgSF molecules that promote metastasis of B16F10 melanoma cells

To identify IgSF molecules that regulate metastasis by modulating tumor-host interactions, we used an experimental lung metastasis model of mouse melanoma B16F10 cells in C57BL/6 mice. We screened for differentially expressed genes between B16F10 and the parental B16 cells (Fig. 1A). We designed PCR primers for 325 murine genes encoding cell-surface IgSF molecules, which were defined as molecules containing Ig-like loops and transmembrane domains or glycosylphosphatidylinositol (GPI) anchor signals (Table S1). RT-PCR of the 325 genes detected 123 genes in B16F10 cells, and subsequent qRT-PCR of the 123 genes in B16F10 and B16 cells showed that the expression levels of 22 genes were over 10-fold higher in B16F10 cells than in B16 cells (Fig. 1B). These genes are considered as potential metastasis-promoting genes and included *Ceacam1*, which promotes melanoma metastasis ^17^ (Table 1). Among these genes, we selected *Cntn1*, *Hfe*, *Igsf3*, *Igsf11*, and *Unc5c* due to their unknown functions in metastasis. Each gene was knocked down in B16F10 cells using two independent shRNAs (Fig. 2A). An experimental lung metastasis model of B16F10 cells demonstrated that the knockdown of *Igsf3* significantly reduced the number of metastatic foci in the lungs after tail vein injection (Fig. 2B– D). A considerable amount of IGSF3 protein expression was detected in B16, and its increased expression in B16F10 cells was confirmed (Fig. 2E). Therefore, we focused on the role of IGSF3 in subsequent studies.

**Figure 1.**
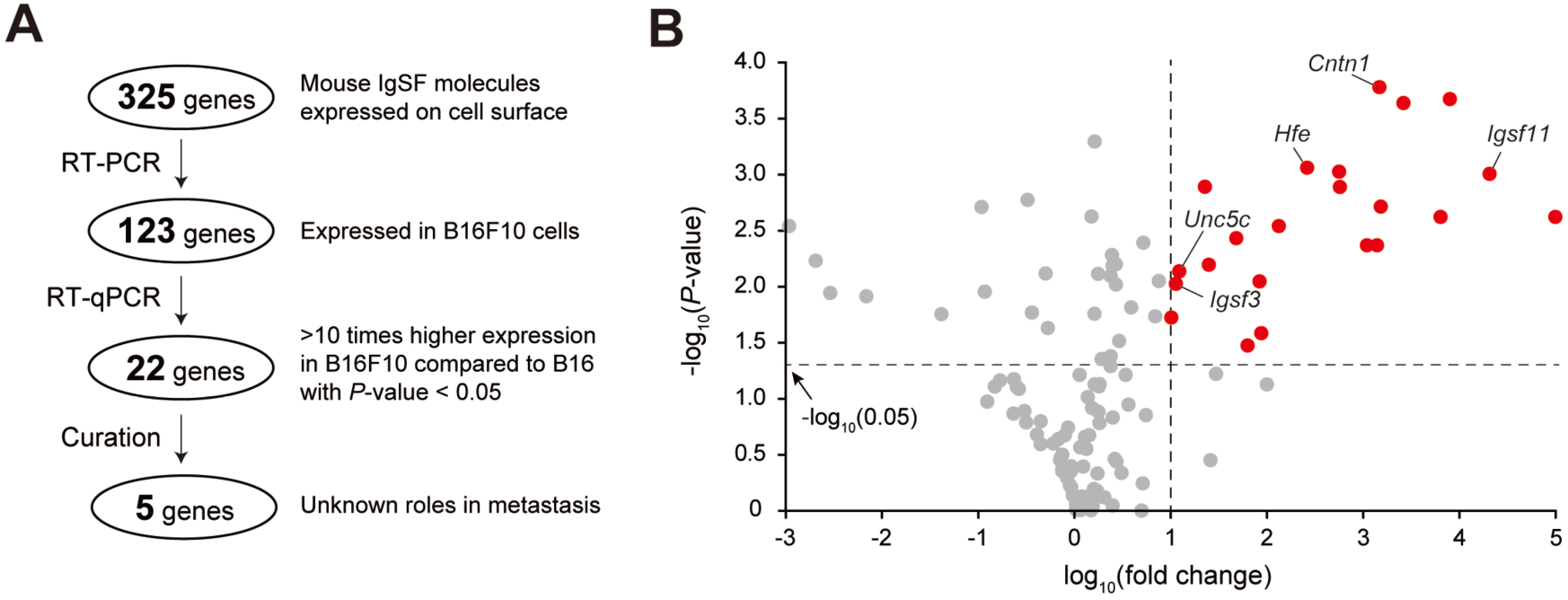
RT-PCR screening for IgSF molecules that potentially promote melanoma metastasis. (**A**) Flowchart illustrating the screening process for genes encoding IgSF molecules that promote metastasis. (**B**) A volcano plot showing the differential expression of genes encoding IgSF molecules in B16F10 cells compared to that in B16 cells. Potential metastasis-promoting genes, defined as those with a fold change > 10 and *p* < 0.05, are highlighted in red.

**Figure 2.**
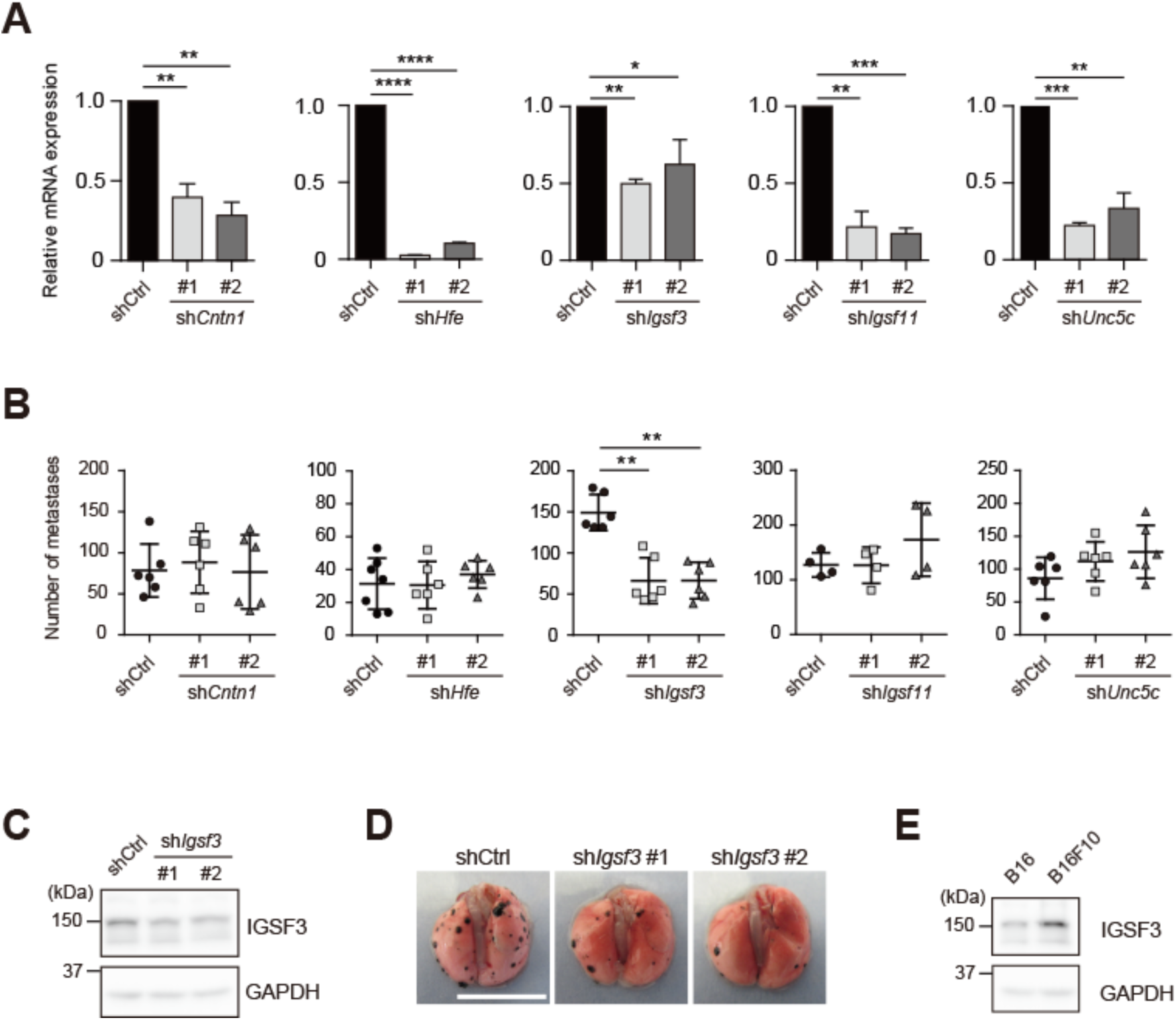
Knockdown of *Igsf3* reduces lung metastasis of B16F10 cells after tail vein injection. (**A**) Each candidate gene was stably knocked down using two independent shRNA vectors. Knockdown efficiency was examined by qRT-PCR. * *p* < 0.05, ** *p* < 0.01, *** *p* < 0.001, **** *p* < 0.0001 by *t*-test (n = 3). (**B**) The number of metastatic foci in the lungs observed two weeks after intravenous injection of B16F10 cells into C57BL/6 mice. ** *p* < 0.01 by Mann– Whitney *U* test. (**C**) Knockdown of IGSF3 protein in B16F10 cells was verified by Western blotting. (**D**) Representative images of the lung after intravenous injection of *Igsf3*-knockdown B16F10 cells. Bar, 1 cm. (**E**) IGSF3 protein expression in B16 and B16F10 was examined by Western blotting.

**Table 1.**
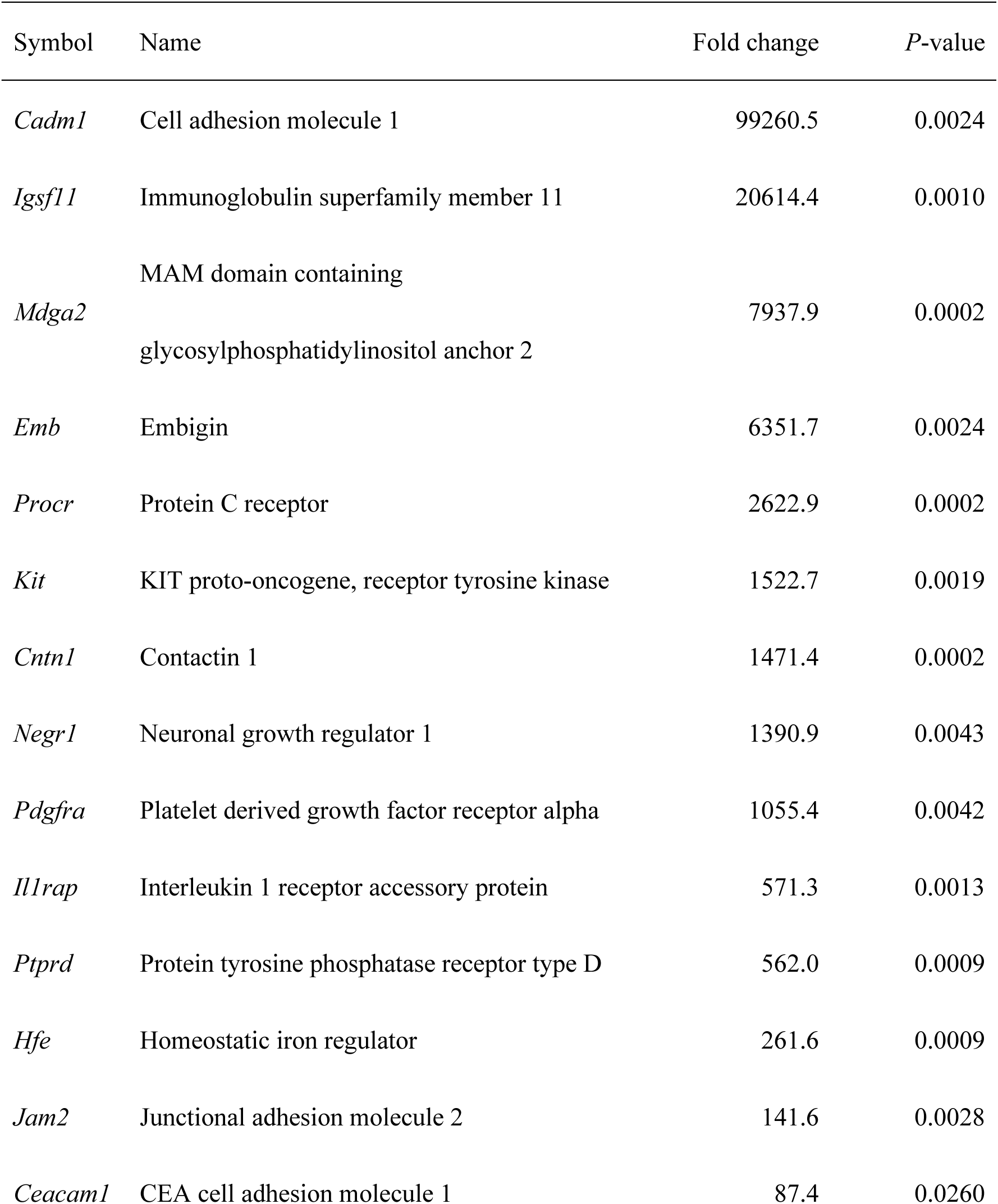
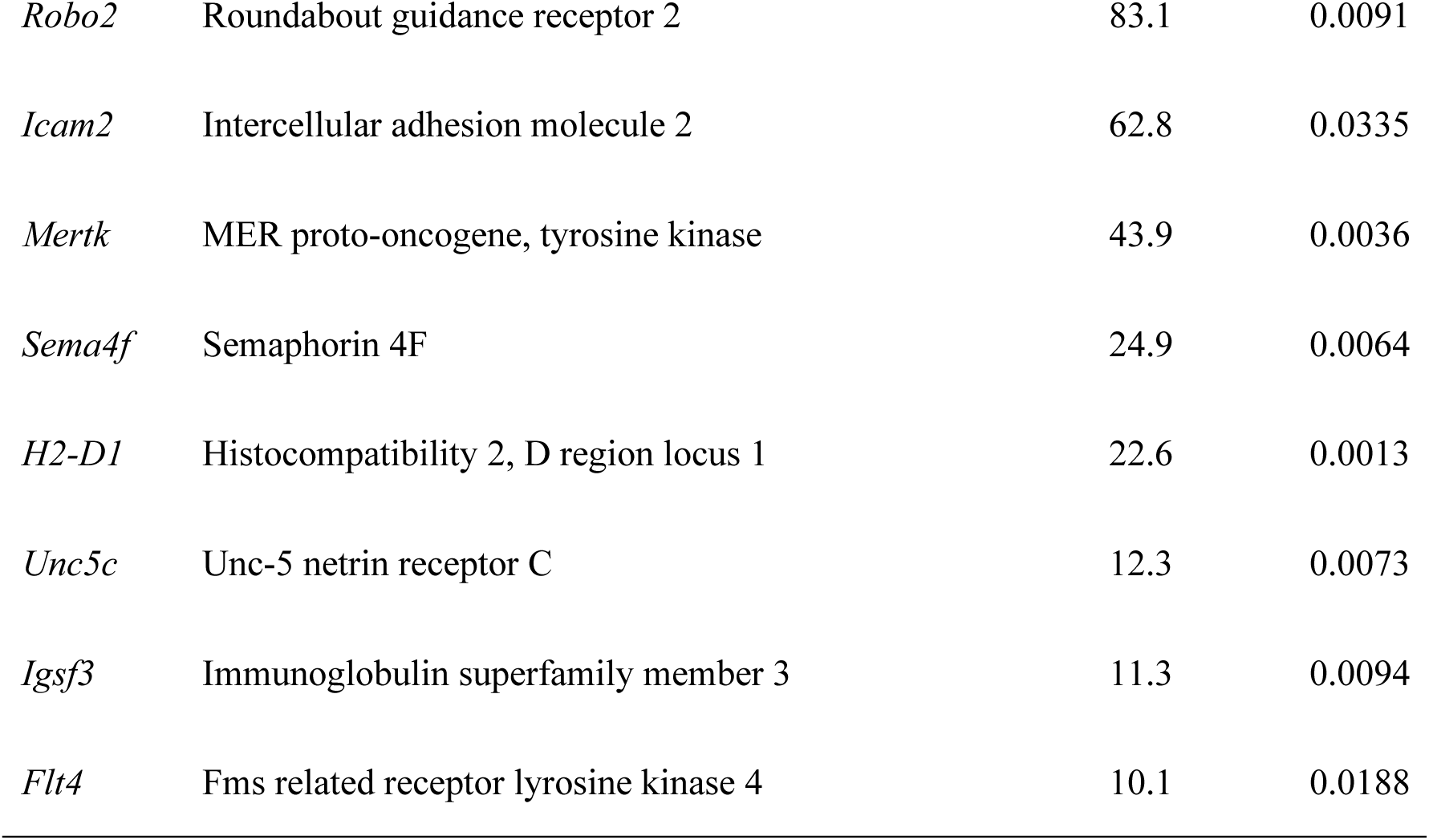
List of IgSF molecules that potentially promote metastasis of B16F10 cells. The table shows the fold change in B16F10 cells compared to that in B16 cells, along with the *p*-values for the potential metastasis-promoting genes (defined as those with fold change > 10 and *p* < 0.05). *P*-values were determined by *t*-test (n = 3).

### IGSF3 promotes adhesion of B16F10 cells to vascular endothelial cells

IGSF3 mRNA is expressed in a wide range of human tissues, with higher expression in the placenta, kidney, and lung ^18^. However, its functions in normal and cancer cells are scarcely understood. To elucidate the mechanism underlying the promotion of metastasis by IGSF3 in B16F10 cells, we examined the involvement of IGSF3 in cell growth using a cell proliferation assay on culture dishes, a colony formation assay in soft agar, and a subcutaneous tumor growth assay in C57BL/6 mice. As shown in Fig. 3A–C, IGSF3 knockdown did not affect the growth of B16F10 cells in any of the assays. Next, we investigated the potential involvement of IGSF3 in cell migration using Transwell and wound-healing assays. As shown in Fig. 3D–E, the knockdown of IGSF3 did not alter cell migration or motility. Then, we hypothesized that IGSF3 might play a role in regulating cancer–stromal interactions. To test this hypothesis, we purified the extracellular domain of IGSF3 fused to an Fc tag (IGSF3-Fc) and examined its binding to stromal cells. We found that IGSF3-Fc bound to the surface of MS1 mouse vascular endothelial cells (vECs) (Fig. 3F). Importantly, the knockdown of IGSF3 in B16F10 cells significantly reduced their adhesion to MS1 cells (Fig. 3G). These results suggest that IGSF3 promotes the metastasis of B16F10 cells by enhancing their adhesion to vECs.

**Figure 3.**
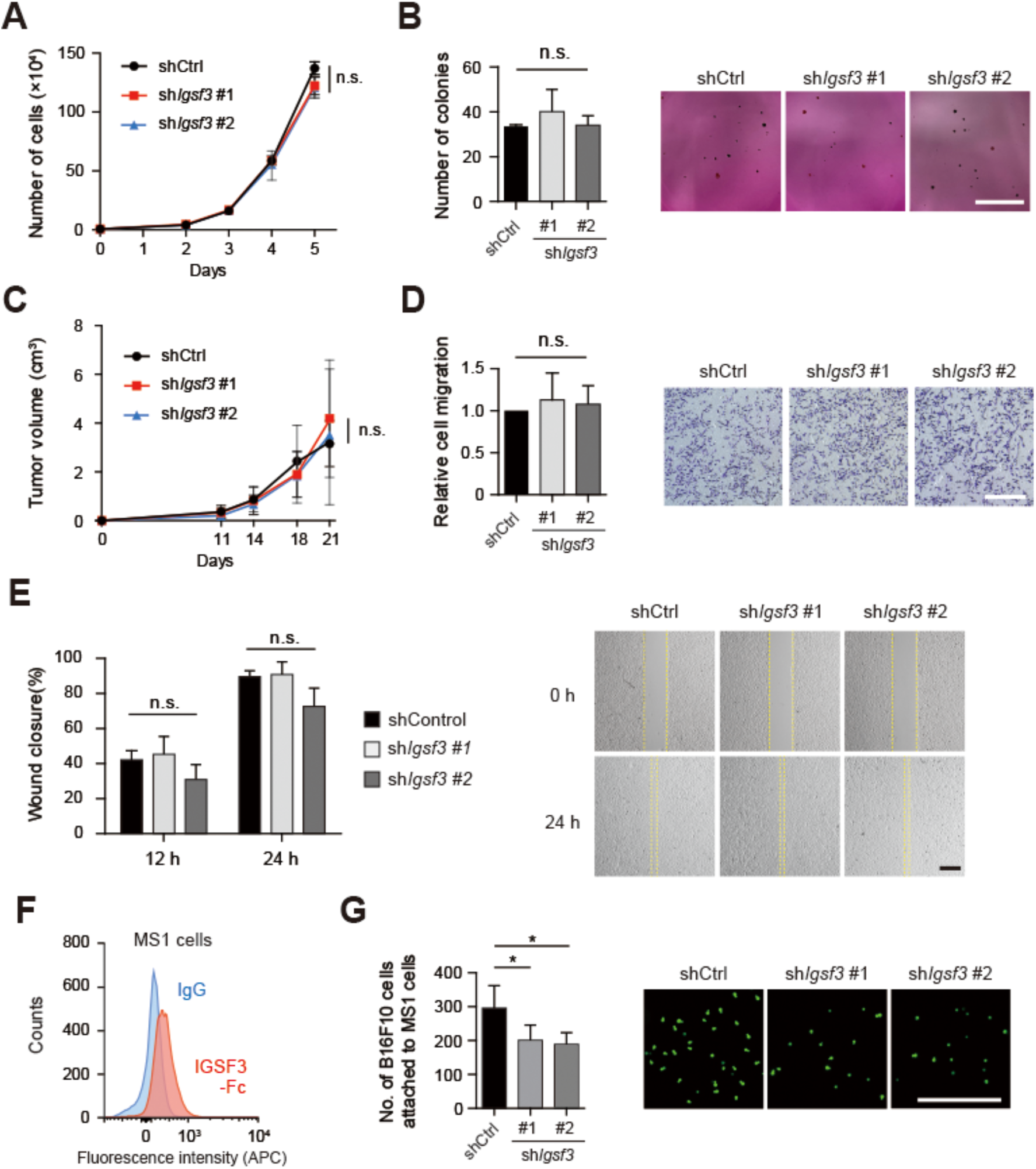
Knockdown of *Igsf3* suppresses adhesion of B16F10 cells to vascular endothelial cells. (**A**) Cell growth assay of B16F10 cells in 6-well plates. (**B**) Soft agar colony formation assay of B16F10 cells. The number of colonies (left) and the representative images of colonies (right) are shown. Bar, 1 cm. (**C**) Subcutaneous tumor growth assay of B16F10 cells in C57BL/6 mice. (**D**) Transwell migration assay of B16F10 cells. The relative number of migrating cells (left) and the representative images of migrating cells (right) are shown. Bar, 500 µm. (**E**) Wound healing assay of B16F10 cells. The area of wound closure at 12 or 24 hours was quantified using the Image J software (left). The wounded area is highlighted in the images on the right. Bar, 500 µm. (**F**) Flow cytometry-based binding analysis of IGSF3-Fc protein to the surface of the murine endothelial cell line, MS1. (**G**) Cell adhesion assay between B16F10 and MS1 cells. The number of calcein AM-labeled B16F10 cells attached to MS1 cells (left) and the representative images of attached cells (right) are shown. Bar, 500 µm. * *p* < 0.05 by *t*-test.

### IGSF3 is a homophilic cell adhesion molecule

We have demonstrated the adhesive activity of IGSF3 between B16F10 and MS1 cells. Then, we examined the binding partner(s) of IGSF3 expressed in MS1 cells. We generated B16 cells stably expressing IGSF3-V5 (Fig. 4A) and analyzed the localization of IGSF3 using immunofluorescence. The IGSF3 signal was found to accumulate at the cell-cell contact site (Fig. 4B), suggesting that IGSF3 mediates adhesion between B16 cells, possibly through the homophilic interactions. To examine whether IGSF3 is involved in a *trans*-homophilic interaction, we co-cultured 293FT cells that had been independently transfected with either IGSF3-V5 or IGSF3-FLAG and then subjected them to immunoprecipitation using an anti-FLAG antibody. A band corresponding to IGSF3-V5 was detected in the precipitates (Fig. 4C), indicating that IGSF3 forms a *trans*-homophilic interaction with neighboring cells. In addition to *trans* interaction, cell adhesion molecules often form *cis* interactions on the same membrane to facilitate mature *trans* interaction ^19^. When 293FT cells are in suspension, the molecular weight of IGSF3 was approximately 160 kDa. In contrast, IGSF3 formed a protein complex of > 500 kDa in a single-cell suspension of 293FT cells in the presence of the crosslinker BS^3^ (Fig. 4D). These results suggest that IGSF3 forms *cis*-homo-multimers or protein complexes with other molecules on the same membrane.

**Figure 4.**
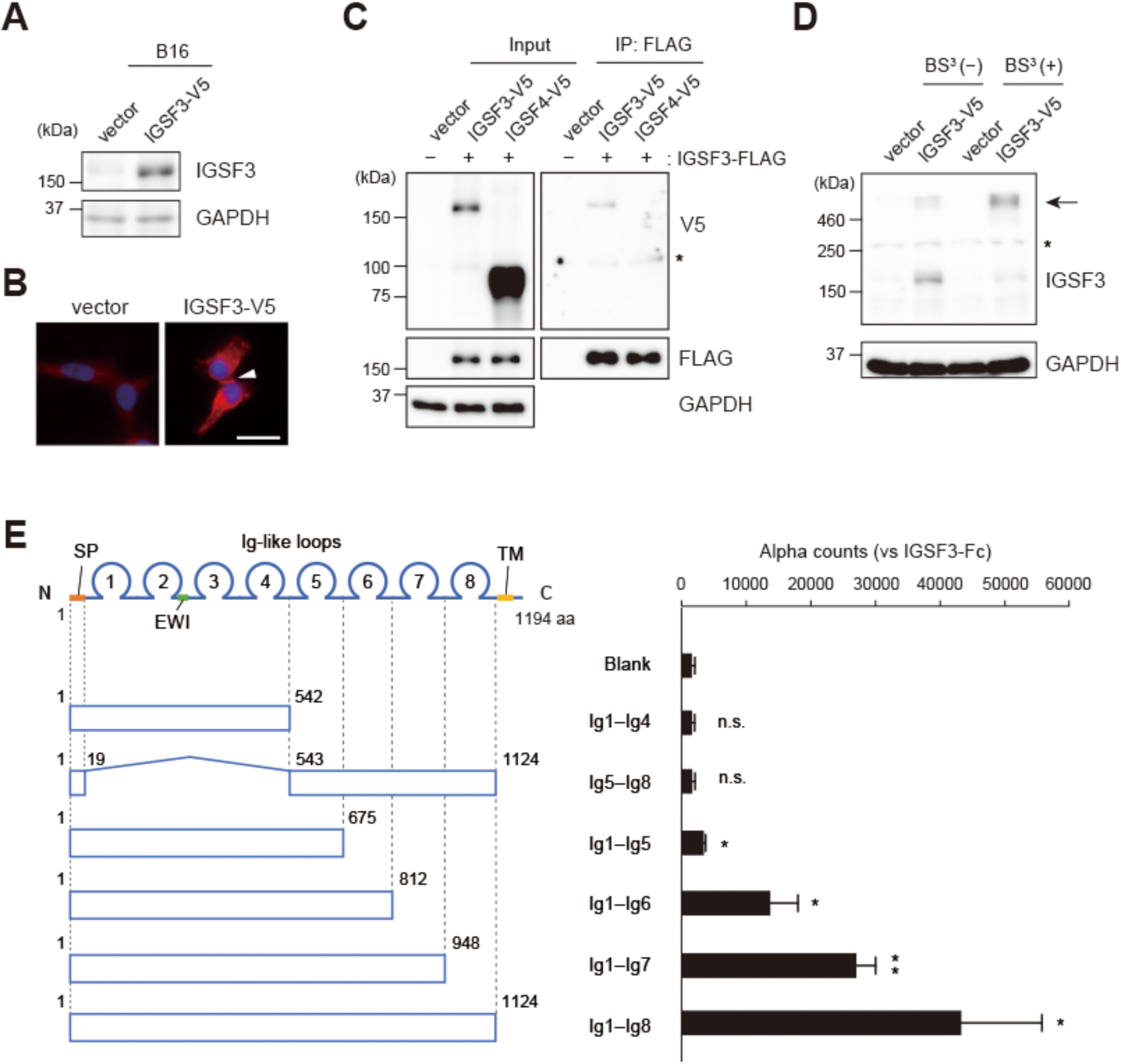
IGSF3 forms *trans*-homophilic interactions between cells. (**A**) Generation of B16 cells expressing IGSF3-V5. (**B**) Recruitment of IGSF3 to the cell-cell contact site in B16 cells, observed by red fluorescence labeled with an anti-V5 antibody (arrowhead). Nuclei were counterstained in blue. Bar, 20 µm. (**C**) The *trans*-homophilic interaction of IGSF3 in the extracellular region was analyzed using an immunoprecipitation assay. IGSF4 served as a negative control. The asterisk denotes non-specific bands. (**D**) *Cis*-multimer formation of IGSF3 was examined by Western blotting in the presence of the crosslinker BS^3^. The putative *cis*-multimer is indicated by the arrow. The asterisk denotes non-specific bands. (**E**) Identification of the domains responsible for the *trans*-homophilic interaction of IGSF3. The binding of a series of truncated IGSF3 proteins to IGSF3-Fc protein was assessed using Alpha technology. SP, signal peptide; EWI, Glu-Trp-Ile motif; TM, transmembrane region. * *p* < 0.05, ** *p* < 0.01 by *t*-test (n = 3, vs Blank).

We also investigated the domain responsible for *trans*-homophilic interaction of IGSF3. We generated a series of truncated mutants of the extracellular domain (ECD) of IGSF3, which were fused to a tandem tag comprising the rat Cd4 domain 3+4 (for stability), FLAG (for detection), and 6 × His (for purification). The binding affinity of the IGSF3 mutant protein to IGSF3-Fc protein was quantified using Alpha technology (Fig. S1). Among the eight Ig-like loops within the IGSF3 ECD, neither the four N-terminal loops (Ig1–Ig4) nor the four C-terminal loops (Ig5– Ig8) bound to IGSF-Fc. In contrast, Ig1–Ig5 showed a significant binding signal, and Ig1–Ig6, Ig1–Ig7, and Ig1–Ig8 yielded even more prominent binding signals (Fig. 4E). These results suggested that the Ig4 and Ig5 loops are necessary for the *trans*-homophilic interaction of IGSF3, whereas Ig6, Ig7, and Ig8 loops likely contribute to stabilizing the interaction.

### The homophilic interaction of IGSF3 promotes extravasation of B16F10 cells

Then, we hypothesized that the homophilic interaction of IGSF3 mediates the adhesion of B16F10 cells to the vECs, given that IGSF3 is reported to be expressed in lung vECs ^20^. MS1 mouse vECs also expressed *Igsf3* mRNA, and introduction of siRNA against *Igsf3* in MS1 markedly reduced its expression (Fig. 5A). The adhesion of B16F10 cells to MS1 cells was suppressed by *Igsf3* knockdown in B16F10 and MS1 cells, whereas *Igsf3* knockdown in both cell types did not synergistically suppress adhesion (Fig. 5B). Moreover, we performed a transendothelial migration assay by seeding B16F10 cells onto an MS1 monolayer in a Transwell chamber (Fig. 5C). Consistent with the results of the cell adhesion assay, *Igsf3* knockdown in B16F10 and MS1 cells reduced the number of migrating B16F10 cells, whereas *Igsf3* knockdown in both cell lines did not synergistically reduce migration (Fig. 5D and 5E). These results suggest that the *trans*-homophilic interaction of IGSF3 from B16F10 cells and vECs promotes adhesion of B16F10 to vECs and their subsequent transendothelial migration. Furthermore, we investigated the role of IGSF3 in the extravasation of B16F10 cells *in vivo*. The localization of B16F10 cells in the lungs 24 h after tail vein injection was determined to be either inside or outside the blood vessels by confocal imaging (Fig. 5F). The results showed that *Igsf3* knockdown reduced the infiltration of B16F10 cells from blood vessels into the lung tissues, from 41.7% to 21.2% of the total cells analyzed (Fig. 5G), suggesting that IGSF3 promotes the extravasation of B16F10 cells into the lung.

**Figure 5.**
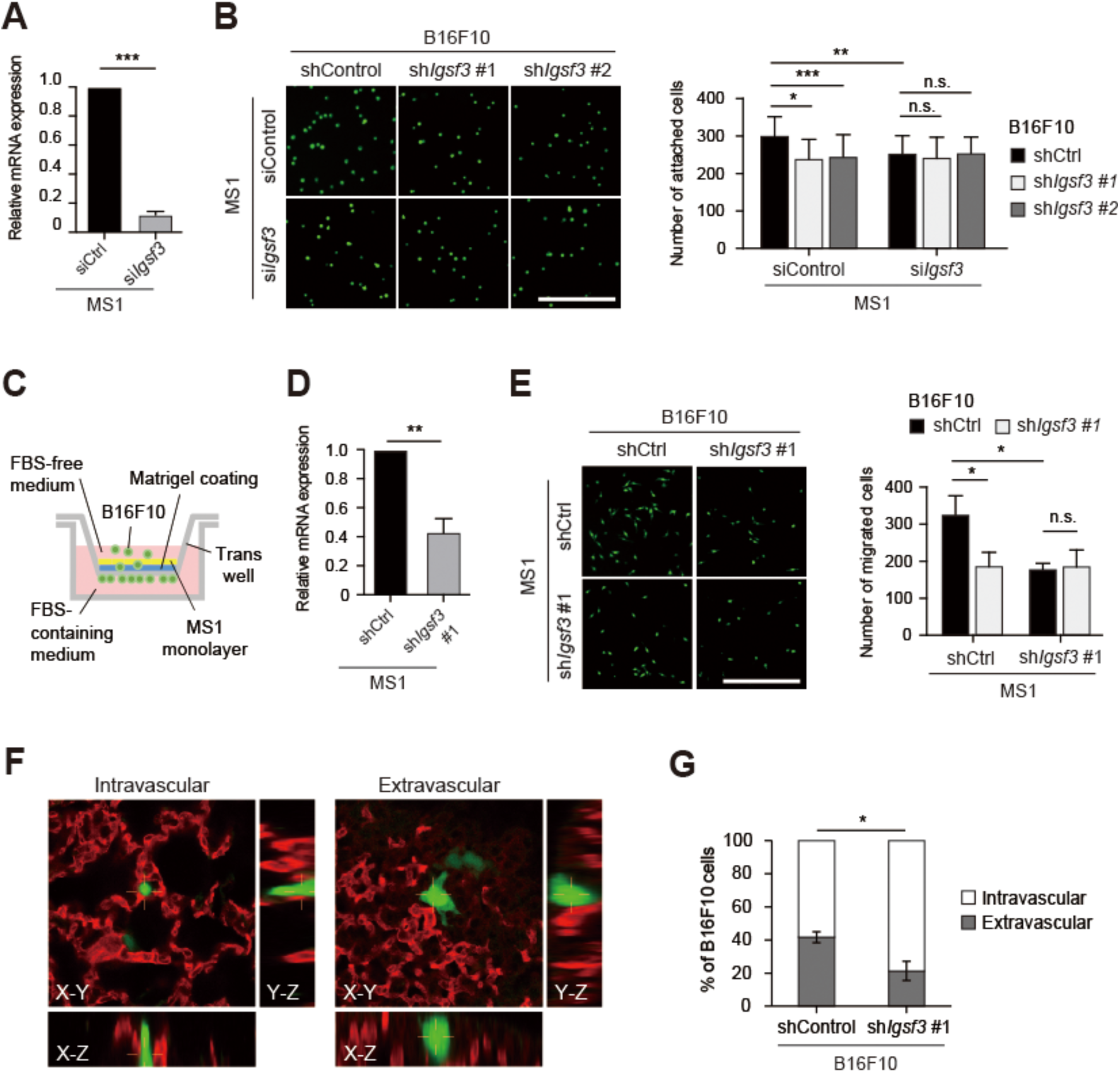
*Trans*-homophilic interaction of IGSF3 mediates the adhesion to the vascular endothelium and subsequent trans-endothelial migration of B16F10 cells. (**A**) Examination of knockdown efficiency of *Igsf3* in MS1 cells using siRNA by qRT-PCR (n = 3, *t*-test). (**B**) Cell adhesion assay. Calcein AM-labeled B16F10 cells with shCtrl or sh*Igsf3* were seeded onto MS1 cells with siCtrl or si*Igsf3*. Representative images (left) and the number of B16F10 cells attached to MS1 cells are shown (right, n = 3, *t*-test). Bar, 500 µm. (**C**) Schematic diagram of the trans-endothelial migration assay. (**D**) Examination of the knockdown efficiency of *Igsf3* in MS1 cells using shRNA by qRT-PCR (n = 3, *t*-test). (**E**) Transendothelial migration assay. Representative images (left) and the number of migrated B16F10 cells are shown (right, n = 3, *t*-test). (**F**) Representative images of B16F10 cells in the lung 24 hours after intravenous injection. Blood vessels were stained using an anti-CD31 antibody (red) to determine whether B16F10 cells expressing EGFP were localized inside (left) or outside (right) of the blood vessels, as observed through confocal microscopy. (**G**) Percentage of intravascular and extravascular B16F10 cells. Approximately 20 cells (ranging from 17 to 24 cells) were analyzed in each group (n = 3, *t*-test). \**p* < 0.05, \*\**p* < 0.01, \*\*\**p* < 0.001.

### The role of IGSF3 in human melanoma

Finally, to understand the role of IGSF3 in human melanoma, we analyzed the expression of IGSF3 in publicly available clinical datasets. The expression of *IGSF3* was significantly higher in primary melanoma tissue than in normal skin tissue in two independent datasets (Fig. 6A). Moreover, higher *IGSF3* expression was correlated with poorer prognosis in melanoma patients in two independent datasets (Fig. 6B). These results suggest that IGSF3 is a potential prognostic factor of human melanoma.

**Figure 6.**
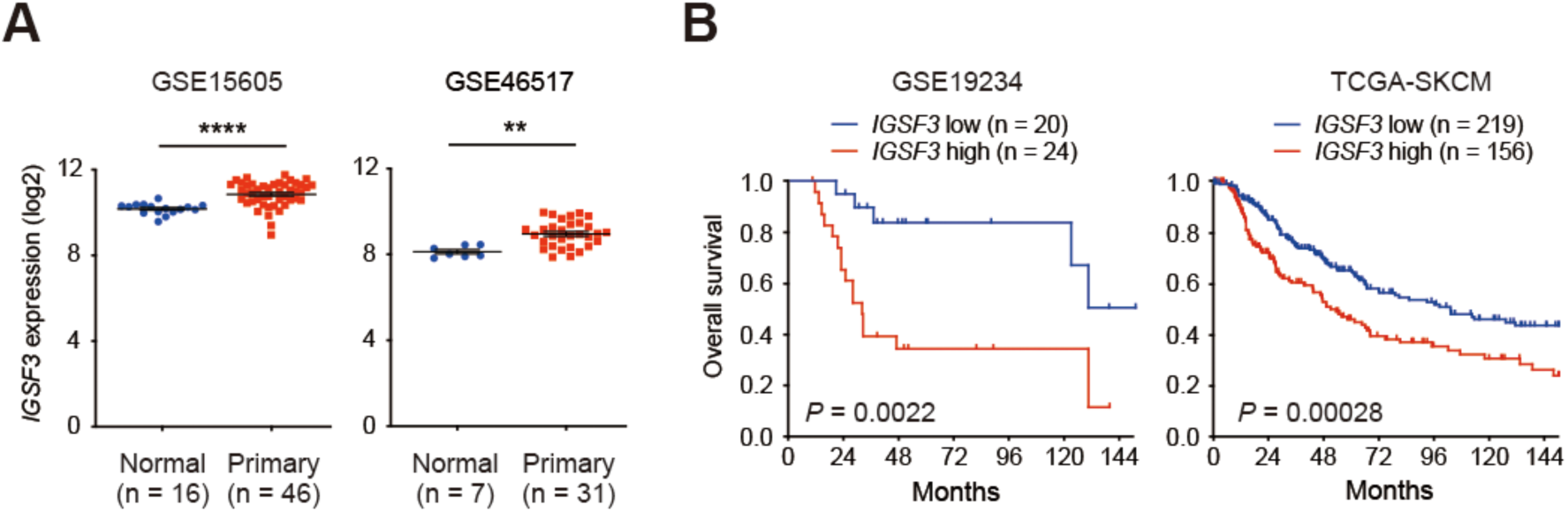
*In silico* analysis of *IGSF3* expression in patients with melanoma. (**A**) Comparison of *IGSF3* expression between human normal skin tissues and primary melanoma using publicly available datasets (GSE15605 and GSE46517). \*\**p* < 0.01 and \*\*\*\**p* < 0.0001 by the Mann– Whitney U test. (**B**) Overall survival analyses of patients with melanoma stratified by high (red) or low (blue) *IGSF3* expression, based on publicly available datasets (GSE19234 and TCGA-SKCM). *P*-values were determined by the log-rank test.

## Discussion

IGSF3 belongs to the EWI subfamily of the IgSF and is characterized by a Glu-Trp-Ile (EWI) motif. IGSF3 forms a complex on the plasma membrane with tetraspanins through this motif and plays critical roles such as maintaining sphingolipid homeostasis during repair after lung injury and regulating cerebellar granule cell differentiation in neuronal morphogenesis ^20, 21^. However, the physiological roles of IGSF3 are not well understood, and its function in mediating intercellular interactions has not yet been recognized. In this study, we demonstrated that IGSF3 acts as a homophilic cell adhesion molecule that mediates the adhesion between melanoma cells and vECs, thereby promoting the extravasation of melanoma cells during metastasis.

Adhesion of cancer cells to vECs is the first step in extravasation. This process requires the expression of ligand–receptor pairs between cancer cells and vECs ^22^. Most cell adhesion molecules, including selectins, integrins, cadherins, and IgSF cell adhesion molecules (IgCAMs), have been implicated in this process. It is generally considered that selectins contributes to the initial attachment, whereas other adhesion molecules, including IgCAMs, facilitate stable adhesion between cancer cells and vECs ^23^. Among the IgCAMs, ICAM-1 and VCAM-1 in vECs are known to interact with integrins in cancer cells ^24^. In addition, homophilic or heterophilic binding of PECAM1, JAM-C, MCAM, AMIGO2, and CADM1 enhances tumor–endothelium adhesion ^11, 25–28^. Our study reveals IGSF3 as another example of such IgCAMs (Fig. 5). Moreover, endothelial IgCAMs serve not only as receptors for their ligands on cancer cells but also trigger a wide range of events in endothelial cells, such as modulation of the cytoskeleton and cell-cell contact. For example, ICAM-1 and VCAM-1 activate Rac1 and reactive oxygen species (ROS)-PKCα pathways, which requires association with tetraspanins for proper transendothelial migration ^29^. Although the cytoplasmic domain of IGSF3 contains only 48 amino acids and lacks conserved functional domains, IGSF3 may trigger downstream signaling in vECs through its association with tetraspanin.

Regarding the molecular basis of the IGSF3 homophilic interaction, IGSF3 has a large ECD consisting of eight Ig-like loops, with Ig4 and Ig5 loops being critical for its *trans* interaction (Fig. 4E). IgCAMs use distinct regions of the protein surface to mediate *trans* interaction. Nectin and Nectin-like/CADM families interact through their Ig1 loop in a handshake-like orientation. In contrast, the L1 family adopts a horseshoe-shaped conformation where the Ig1 and Ig2 loops fold back to interact with lg4 and Ig3 loops, respectively. These horseshoe-shaped monomers facilitate *trans*-homophilic interaction ^19, 30^. The requirement of the Ig4 and Ig5 loops for *trans*-homophilic interaction of IGSF3 suggests that its ECD may have a curved structure, which presents an interaction interface within the Ig4 and Ig5 loops, similar to the L1 family. Furthermore, while IgCAMs often form *cis* homodimers that precede *trans* interactions, IGSF3 forms *cis*-homo-multimers or stable complexes with other molecules (Fig. 4D). Such large complex formation on the same membrane may contribute to the function of IGSF3. Therefore, crystal structure analysis of IGSF3 is expected to elucidate its unique *cis* and *trans* interaction formations.

IGSF3 promotes the growth and migration of hepatocellular carcinoma cells by activating the NF-κB pathway ^31^ and drives glioma progression through its association with Kir4.1, leading to potassium dysregulation ^32^. In human cancer, elevated IGSF3 expression correlates with a worse prognosis in endometrial and pancreatic cancers; in contrast, it is associated with a better prognosis in renal cancer ^33^. Our study demonstrated that IGSF3 is not directly involved in the proliferation or migration of melanoma cells (Fig. 3). However, it promotes metastasis by enhancing the adhesion of melanoma cells to vECs and subsequent transendothelial migration (Fig. 5). These observations suggest that the role of IGSF3 in cancer is context-dependent. Given its ubiquitous expression and low tissue- or cell-type specificity, the context-dependent roles of IGSF3 could be modulated by its associated molecules, such as Kir4.1, which is predominantly expressed in neural cells. Further studies are required to understand the oncogenic and tumor-suppressive roles of IGSF3, particularly its *cis*-interacting molecules and downstream cascades.

To the best of our knowledge, this is the first report of the involvement of IGSF3 in cancer metastasis. IGSF3 drives lung metastasis by fostering the preferential adhesion of melanoma cells to vECs through *trans*-homophilic interactions, which offers a potential prognostic factor for melanoma. Thus, targeting the homophilic interaction of IGSF3 may provide a therapeutic strategy for combating the metastatic spread of melanoma, and possibly other cancers.

## Acknowledgements

This study was supported in part by AMED under Grant Number JP22ama221307, JSPS KAKENHI Grant Number 20H05028 and 21K07091 to TI, 20H03525 and 20K21539 to YM.

## References

1 Lambert AW, Pattabiraman DR, Weinberg RA. Emerging Biological Principles of Metastasis. Cell. 2017; 168: 670–691.

2 McAllister SS, Weinberg RA. The tumour-induced systemic environment as a critical regulator of cancer progression and metastasis. Nat Cell Biol. 2014; 16: 717–727.

3 van der Weyden L, Arends MJ, Campbell AD, et al. Genome-wide in vivo screen identifies novel host regulators of metastatic colonization. Nature. 2017; 541: 233–236.

4 van der Weyden L, Harle V, Turner G, et al. CRISPR activation screen in mice identifies novel membrane proteins enhancing pulmonary metastatic colonisation. Commun Biol. 2021; 4: 395.

5 Zinn K, Özkan E. Neural immunoglobulin superfamily interaction networks. Curr Opin Neurobiol. 2017; 45: 99–105.

6 Barclay AN. Membrane proteins with immunoglobulin-like domains--a master superfamily of interaction molecules. Semin Immunol. 2003; 15: 215–223.

7 van Elsas A, Hurwitz AA, Allison JP. Combination immunotherapy of B16 melanoma using anti-cytotoxic T lymphocyte-associated antigen 4 (CTLA-4) and granulocyte/macrophage colony-stimulating factor (GM-CSF)-producing vaccines induces rejection of subcutaneous and metastatic tumors accompanied by autoimmune depigmentation. J Exp Med. 1999; 190: 355–366.

8 Chen Q, Massagué J. Molecular pathways: VCAM-1 as a potential therapeutic target in metastasis. Clin Cancer Res. 2012; 18: 5520–5525.

9 Sarbassov DD, Guertin DA, Ali SM, Sabatini DM. Phosphorylation and regulation of Akt/PKB by the rictor-mTOR complex. Science. 2005; 307: 1098–1101.

10 Ito T, Nakamura A, Tanaka I, et al. CADM1 associates with Hippo pathway core kinases; membranous co-expression of CADM1 and LATS2 in lung tumors predicts good prognosis. Cancer Sci. 2019; 110: 2284–2295.

11 Kasai Y, Gan SP, Funaki T, et al. Trans-homophilic interaction of CADM1 promotes organ infiltration of T-cell lymphoma by adhesion to vascular endothelium. Cancer Sci. 2022; 113: 1669–1678.

12 Ito T, Kasai Y, Kumagai Y, et al. Quantitative Analysis of Interaction Between CADM1 and Its Binding Cell-Surface Proteins Using Surface Plasmon Resonance Imaging. Front Cell Dev Biol. 2018; 6: 86.

13 Raskin L, Fullen DR, Giordano TJ, et al. Transcriptome profiling identifies HMGA2 as a biomarker of melanoma progression and prognosis. J Invest Dermatol. 2013; 133: 2585–2592.

14 Kabbarah O, Nogueira C, Feng B, et al. Integrative genome comparison of primary and metastatic melanomas. PLoS One. 2010; 5: e10770.

15 Bogunovic D, O’Neill DW, Belitskaya-Levy I, et al. Immune profile and mitotic index of metastatic melanoma lesions enhance clinical staging in predicting patient survival. Proc Natl Acad Sci U S A. 2009; 106: 20429–20434.

16. Genomic Classification of Cutaneous Melanoma. Cell. 2015; 161: 1681–1696.

17 Wicklein D, Otto B, Suling A, et al. CEACAM1 promotes melanoma metastasis and is involved in the regulation of the EMT associated gene network in melanoma cells. Sci Rep. 2018; 8: 11893.

18 Saupe S, Roizès G, Peter M, Boyle S, Gardiner K, De Sario A. Molecular cloning of a human cDNA IGSF3 encoding an immunoglobulin-like membrane protein: expression and mapping to chromosome band 1p13. Genomics. 1998; 52: 305–311.

19 Honig B, Shapiro L. Adhesion Protein Structure, Molecular Affinities, and Principles of Cell-Cell Recognition. Cell. 2020; 181: 520–535.

20. Schweitzer KS, Jinawath N, Yonescu R, et al. IGSF3 mutation identified in patient with severe COPD alters cell function and motility. JCI Insight. 2020; 5.

21 Usardi A, Iyer K, Sigoillot SM, Dusonchet A, Selimi F. The immunoglobulin-like superfamily member IGSF3 is a developmentally regulated protein that controls neuronal morphogenesis. Dev Neurobiol. 2017; 77: 75–92.

22 Witz IP. The selectin-selectin ligand axis in tumor progression. Cancer Metastasis Rev. 2008; 27: 19–30.

23 Reymond N, d’Água BB, Ridley AJ. Crossing the endothelial barrier during metastasis. Nat Rev Cancer. 2013; 13: 858–870.

24 Madsen CD, Sahai E. Cancer dissemination--lessons from leukocytes. Dev Cell. 2010; 19: 13–26.

25 Kanda Y, Osaki M, Onuma K, et al. Amigo2-upregulation in Tumour Cells Facilitates Their Attachment to Liver Endothelial Cells Resulting in Liver Metastases. Sci Rep. 2017; 7: 43567.

26 Jouve N, Bachelier R, Despoix N, et al. CD146 mediates VEGF-induced melanoma cell extravasation through FAK activation. Int J Cancer. 2015; 137: 50–60.

27 Langer HF, Orlova VV, Xie C, et al. A novel function of junctional adhesion molecule-C in mediating melanoma cell metastasis. Cancer Res. 2011; 71: 4096–4105.

28 DeLisser H, Liu Y, Desprez PY, et al. Vascular endothelial platelet endothelial cell adhesion molecule 1 (PECAM-1) regulates advanced metastatic progression. Proc Natl Acad Sci U S A. 2010; 107: 18616–18621.

29 van Buul JD, Kanters E, Hordijk PL. Endothelial signaling by Ig-like cell adhesion molecules. Arterioscler Thromb Vasc Biol. 2007; 27: 1870–1876.

30 Wei CH, Ryu SE. Homophilic interaction of the L1 family of cell adhesion molecules. Exp Mol Med. 2012; 44: 413–423.

31 Sheng P, Zhu H, Zhang W, et al. The immunoglobulin superfamily member 3 (IGSF3) promotes hepatocellular carcinoma progression through activation of the NF-κB pathway. Ann Transl Med. 2020; 8: 378.

32 Curry RN, Aiba I, Meyer J, et al. Glioma epileptiform activity and progression are driven by IGSF3-mediated potassium dysregulation. Neuron. 2023; 111: 682–695.e689.

33 Ding Y, Chen J, Li S, et al. EWI2 and its relatives in Tetraspanin-enriched membrane domains regulate malignancy. Oncogene. 2023; 42: 861–868.

