## Supplementary material for "IGSF3 is a homophilic cell adhesion molecule that drives lung metastasis of melanoma by promoting adhesion to vascular endothelium": Document S1

**Document S1. Extended Materials and Methods**

**Cell lines**

Mouse melanoma cell lines B16 and B16F10 were obtained from the Cell Resource Center for Biomedical Research (Sendai, Japan). The mouse vascular endothelial cell line MS1 was purchased from the American Type Culture Collection (Manassas, VA, USA). The human embryonic kidney cell lines 293FT and Expi293-F were purchased from Thermo Fisher Scientific (Waltham, MA, USA). B16, B16F10, MS1, and 293FT cells were cultured in DMEM (Nacalai Tesque, Kyoto, Japan) supplemented with 10% fetal bovine serum (Biowest, Nuaille, France), 100 units/mL penicillin, and 100 μg/mL streptomycin (Sigma-Aldrich, St. Louis, MO, USA) in a humidified incubator at 37°C with 5% $CO_2$ (Panasonic Healthcare, Tokyo, Japan). Expi293-F cells were cultured in Expi293 Expression Medium (Thermo Fisher Scientific) supplemented with 50 units/mL penicillin and 50 μg/mL streptomycin. The cultures were shaken at 125 rpm on a rotary shaker (Wakenbtech, Kyoto, Japan) in an incubator at 37°C with 8% $CO_2$.

**Antibodies**

Mouse monoclonal anti-IGSF3 (503621), anti-FLAG (M2), anti-GAPDH (6C5), and rabbit polyclonal anti-V5 (PM003) antibodies were purchased from Thermo Fisher Scientific, Sigma-Aldrich, Merck Millipore (Darmstadt, Germany), and MBL (Nagoya, Japan), respectively. Secondary antibodies conjugated with horseradish peroxidase (HRP) and Alexa Fluor 546 were obtained from Merck Millipore and Thermo Fisher Scientific, respectively.

**Western blotting**

Cell lysates were extracted using a lysis buffer [50 mM Tris-HCl (pH 7.5), 150 mM NaCl, 1 mM EDTA, 1% Triton X-100] containing a protease inhibitor cocktail (200 μM AEBSF, 10 μM leupeptin, 1 μM pepstatin A). Protein samples were fractionated by 7.5% SDS-PAGE and transferred to polyvinylidene difluoride membranes (Merck Millipore). Membranes were incubated with primary antibodies, and their binding was detected using HRP-conjugated secondary antibodies and Pierce Western Blotting Substrate (Thermo Fisher Scientific). Signals were visualized using Amersham Imager 680 (GE Healthcare, Little Chalfont, UK).

**Immunofluorescence**

The cells were seeded onto glass coverslips coated with poly-L-lysine. After two days, the cells were fixed with 4% PFA, permeabilized with 0.2% TritonX-100, and blocked with 2% normal goat serum. The cells were labeled with an anti-V5 primary antibody and an Alexa Fluor 546-conjugated secondary antibody. The nuclei were counterstained with Hoechst 33342. Cells were imaged using an epifluorescence microscope (Axio Observer D1; Carl Zeiss, Oberkochen, Germany).

**Production of IGSF3 proteins**

The Gateway-based destination vector, pHEK-Cd4-FLAG-His, was generated by inserting a DNA fragment containing a Gateway cassette, followed by a tandem tag composed of rat Cd4 domain 3+4 (aa 209–302), FLAG, and 6 × His into the *Sma*I-*Sph*I site of the pHEK293 Ultra Expression Vector II (Takara Bio, Shiga, Japan). The expression vectors for IGSF3 ECD and its mutants with a Cd4-FLAG-His tag were generated using Gateway recombination with LR clonase between the pENTR/D-TOPO and pHEK-Cd4-FLAG-His vectors. The expression vector for IGSF3 ECD with a human $IgG_2$-Fc tag (IGSF3-Fc) was obtained by Gateway recombination between the pENTR/D-TOPO and pHEK-Fc vectors, as previously described [1]. The IGSF3-Cd4-FLAG-His and IGSF3-Fc proteins were produced according to a method described previously [2]. Briefly, the expression vectors and the pHEK293 Enhancer Vector (Takara Bio) were transiently transfected into Expi293-F cells using ScreenFect UP-293 (Fujifilm Wako Pure Chemical, Osaka, Japan). After culturing for five days on a shaker, the supernatant was filtered and purified using Ni or Protein A Sepharose (GE Healthcare). The obtained proteins were checked for their purity by SDS-PAGE followed by silver staining using the Silver Stain MS Kit (Fujifilm Wako Pure Chemical) and stored at −80°C until further use.

**References**


1 Ito, T. *et al.* Quantitative Analysis of Interaction Between CADM1 and Its Binding Cell-Surface Proteins Using Surface Plasmon Resonance Imaging. *Front Cell Dev Biol* **6**, 86, doi:10.3389/fcell.2018.00086 (2018).

2 Kasai, Y. *et al.* Trans-homophilic interaction of CADM1 promotes organ infiltration of T-cell lymphoma by adhesion to vascular endothelium. *Cancer Sci* **113**, 1669-1678, doi:10.1111/cas.15307 (2022).
