## Supplementary material for "IGSF3 is a homophilic cell adhesion molecule that drives lung metastasis of melanoma by promoting adhesion to vascular endothelium": Table S1

**Supplementary Table S1.** List of primer sets for detecting the expression of genes encoding cell-surface IgSF in mice.

| Gene | Sense primer (5' → 3') | Antisense primer (5' → 3') | Amplicon |
|---|---|---|---|
| *Adgra2* | TCAACCACAGCTCCATCCAC | TGGTAGTTGGTGAGCGTGAC | 118 (bp) |
| *Adgra3* | GTGTACTCCACGTACTCGCC | TCCCTTCGCCCATAGTCAGA | 140 |
| *Adgra5* | CCGATTGTCTGTGAGGTCCC | TCTGAGTGGCTTCGCCTGTC | 140 |
| *Ager* | GGTTCTTGCTCTATGGGGAGC | GAGAGGACCTTCCAAGCTTCA | 159 |
| *Alcam* | TCTGCGATAAGTATTCCAGAGCAC | CAGCCAGTAGACGACACCAG | 147 |
| *Amigo1* | ATCCTCGCCGAGCAGAACAT | GCCCTGGCTACCTCAAAGAG | 156 |
| *Amigo2* | TAGACCGACGGCTGGCTAAG | TGCCTTAACTCCCTCCACCG | 90 |
| *Amigo3* | GTGGAGTCTGGTGGGCATAC | CTGCTGGGTTGTGAGCCTTA | 103 |
| *Axl* | CATACATCGGGACCTGGCTG | CCTTGGCGGTAGTAATCCCC | 114 |
| *Bcam* | ACTGTGCTGCTCACTACGAC | GTTCCGTGGGATAGTGCAGG | 90 |
| *Boc* | AGTGACGCTAGCCAATCTCC | CTCCAGCCACTCCTGTTTGA | 151 |
| *Bsg* | GGCACCATCCAAACCTCTGT | GCGGTCATCTGCGTCCACTA | 174 |
| *Btla* | TGCTTGGGACTCCTCGGTTA | GCACTGGACACTCTTCATCATT | 108 |
| *Btn1a1* | ACGTCAGAGTCCAAGAAGCAT | AGGCCAGTAAGATGATAGCCA | 208 |
| *Btn2a2* | TGGAGACGAACCCTCTTACATG | CACATGGACGGCAGTCAAATC | 157 |
| *Btnl2* | GACCGAACTGGCTTCCGTTA | GGTAATAGCGGGATCGGAGC | 146 |
| *Btnl9* | GGGAAGCGTGAACCTACACA | GAGCCCGATCCTGCTACTTC | 126 |
| *Cadm1* | ATTCTGGGCCGCTATTTTG | TGTCCTCCTTCTGCATTGATT | 113 |
| *Cadm2* | GACAGTACACCTGCTCCTTATT | AATCCACTAATCTGAGGCTTCTC | 97 |
| *Cadm3* | GAATCTCTGAAGGGAGCCGAC | CCACGCCCCTCACAATGTAAC | 134 |
| *Cadm4* | CATCGGTTCCTTACGCCATTG | CTTCTCCCTGCTCATCCAAGC | 150 |
| *Cd1d1* | GGTCTCCTAGAGGCAGGGAA | GCCAGAGACATGACACACCA | 117 |
| *Cd2* | GCACCATTCAAGTGTGAGGC | CCAAGAGCACCAAGAGGAGT | 136 |
| *Cd4* | CACTCAAGGGAAGACGCTGG | CGATCAAACTGCGAAGGCG | 177 |
| *Cd7* | CAGGAAAGTCAGTGCCCGT | CCGGTGAAGAAGAAGCCTACA | 141 |
| *Cd8a* | CGTGCCAGTCCTTCAGAAAGT | CAATCTTCTGGTCTCTGGGGC | 108 |
| *Cd8b1* | CGCTGATCATTTGTGAAACTGTTT | GAGTGGCCGTCTACTTTTACTGTGT | 77 |
| *Cd19* | ACCAGTTGGCAGGATGATGG | GCTGAGGAGCTGCATAGAGG | 128 |
| *Cd22* | AGCACTCCTCCAAAGTGCTC | CCACCTCTGTGGGATTGACC | 93 |
| *Cd28* | ACCCTCCGCCTTACCTAGAC | AAAACAGGACTCCAGCAACCA | 132 |
| *Cd33* | CCGCTGTTCTTGCTGTGTG | AAGTGAGCTTAATGGAGGGGTA | 139 |

| | | | |
|---|---|---|---|
| *Cd47* | CACGGCCCCCTTTTGATTTC | GTCCGTCACTTCCCTTCACC | 135 |
| *Cd48* | CTGCGTGAAACTGAGAACGAG | TGCTGGTCCTTTACCTCACA | 140 |
| *Cd79a* | CAGGGACGCTGCTGCTATTC | GTCATCAAGGTTCAGGCCCTC | 110 |
| *Cd79b* | CCAGCAATGACAAGCAGTGAC | TCCGCTTTTTGGCTGCAAAC | 94 |
| *Cd80* | CCCCAGAAGACCCTCCTGATAG | CCGAAGGTAAGGCTGTTGTTTG | 172 |
| *Cd83* | CCCCAAGGAAGCTACAGAGTCAA | GGGGAGGTGACTGGAAGAAAAG | 183 |
| *Cd84* | AAAGCAGCTACCACACCAGT | ATTACCACCGGGTCTGCATC | 140 |
| *Cd86* | GCAGCACGGACTTGAACAAC | TTGTAAATGGGCACGGCAGA | 194 |
| *Cd96* | TCCACCTAGGTTCACCTTTTCA | GGAATTGTTGACGTGGTGATGG | 88 |
| *Cd101* | ACTCGGTCCTGCTGCATATC | CAGACAGGGTGTCGGGAAC | 141 |
| *Cd160* | AGCCATATCAACGGCACTCTC | TTTATGGACCCGGCTTCTCTG | 178 |
| *Cd200* | AGGAACAGCTTGCCTTACCC | TCCTGTCCCAGTACCCTTCC | 139 |
| *Cd200r1* | CAAATTGCCAAAATTAGAAGCTACTTC | CACAGTATCATAGAGTGGATTGCTCTT | 100 |
| *Cd200r2* | CTGCTCGTTGTCATTGGTTTG | TTATCTCCAGTCTGCCTCCA | 76 |
| *Cd226* | AGTTGGTTCCCCAAAAGAGG | AGCCATCTCTGGAAACAGTTCA | 136 |
| *Cd244* | TCAAGGACTCACGAGCCAG | CTATCTCCAGGGAAAGTCTGCT | 157 |
| *Cd274* | TAGTGTCCACGGTCCTCCTC | ACACTGCTTACGTCTCCTCG | 126 |
| *Cd276* | TCTTCCCTGACCTGTTGGTG | CATGCTGGGCTTCGAGTAGG | 158 |
| *Cd300a* | CAGGACCAACACTAGAGACACC | AGGTCCCCACCAACAGAAAC | 168 |
| *Cd300c* | GCTGTGCTACTCTCCTTGGT | CTGAGGCACTGGTCTTGACA | 199 |
| *Cd300e* | GAGGATGAGGCTATGTGCAGG | ATGTCGTGTCATGCGGTCC | 176 |
| *Cd300lb* | GGTATCCCGCTGAGATTTGGA | TCTTGTTGGTTTGCCATCTTGA | 211 |
| *Cd300ld* | TACAACCAAGCGCTGAGAACA | AGCAGGGGTAACTCCACAAAG | 136 |
| *Cd300lf* | ATTTGTCATTGCTGGTCCCCT | GACACACGGTTCTTCTTCACC | 241 |
| *Cd300lg* | GGCCCCAGTCCTGATACTC | ACAGCAGAAGGGTTGAGGTTTC | 228 |
| *Cdon* | TCTGCACACACAAACTCCCTG | GAAGGGGACTGTTGGTTTTGG | 175 |
| *Ceacam1* | TCGAATCCAGTCAGCGTCAG | CCAGCCCTGCTATTAGAGCC | 142 |
| *Ceacam3* | ATAGCAGAGGTGTGACGGGA | GGCGTCCACAGGTCAAAATG | 195 |
| *Ceacam19* | AACCCTGCCTGGACAATGAA | CATGGACAGGGAGGAGTGAG | 82 |
| *Ceacam20* | GTCCAACCCTGTCACCAACT | CCAAGGTTGGGAACTCGATA | 95 |
| *Chl1* | ACAGATCGGATGTCAAGAGGC | ACGGTGGTCTGCCTGGATTA | 82 |
| *Clmp* | CTCACAGAAGGTCCGTCGGC | GTGTGAGTTCCCAGCGTTCC | 143 |
| *Cntfr* | ACGCAGAAACACAGTCCACA | TCTGTCCCGTTTACCCTCCA | 125 |
| *Cntn1* | GGGTCCATTTATGCCAACGC | AAACTTGGGTTTTGGTGCGG | 147 |

| | | | |
|---|---|---|---|
| *Cntn2* | GGAGCCTGTGCTACAAGACA | CGGCTTTCCAGTAGCGAAT | 65 |
| *Cntn3* | CCTGCTTTCATTCATTGGCTG | GGGTTGCCTCTTGCTTCACA | 141 |
| *Cntn4* | CCCGGCTGAGAAAGGAACAA | GCCATCAGCTCTTCGCCATA | 88 |
| *Cntn5* | CAGCAACGTGAGTGGAAGAA | CCTCAAAGGGTGTGAGAGGA | 215 |
| *Cntn6* | ATACGGCTTGGTTCTGCCTG | TCAGGGAAATGTCTCGGGAT | 98 |
| *Crtam* | AGCTCCAAATACCAGCTTCTTC | GTCACTCTCACTTGCTTCGTC | 137 |
| *Csf1r* | GCCTCTTCCTCTGTTCCCTTTC | GCTAGTTCTGTGAGGACGGG | 117 |
| *Csf3r* | TTAACGACGGGGCTAGAAAG | AGGGGCTTAACAATACCACTC | 147 |
| *Ctla4* | ATGGCTTGTCTTGGACTCCG | CACCACTGAAGGTTGGGTCA | 138 |
| *Cxadr* | CTGTGCTTCGTGCTCTTGTG | TTCGATCCTCTGTTCGGGTG | 78 |
| *Dcc* | GCTGAAACAGTGCGTGTGG | AATAATCAACGGGGTCAGTGGG | 175 |
| *Dscam* | TGCTCCCATCCCTACCCTAC | ACGCTATACTGCCGATTCCG | 101 |
| *Dscaml1* | AGGGCCTCTTTATGGCATGT | CCCACAGAACTGGAGAAGGTC | 141 |
| *Emb* | GGGGGATTCTACTGTGCTGA | GAGTGAGCGTCAATGGGAAC | 105 |
| *Ermap* | CTCTTCACCATAGCTGTCTTCC | CAAGTTGGCTGTGTCCCTAA | 93 |
| *Esam* | CCCAGGTCTTCTTTGGACCAG | CCACTCTGTTTTGAGCCTTGC | 111 |
| *F11r* | CTGTAATGGGCACCGAGGG | CCGAACCCTTGCCTTGTACC | 93 |
| *Fam187a* | GCCTACCTGGCTGATATGAGC | CAACCGCCCATCAAAGTCTGT | 135 |
| *Fcamr* | CTCCCTTTCAGGTACAAATGCA | TCTTTGATGCCTGTTGACTGAG | 115 |
| *Fcer1a* | GTCTCCATTAGAGAGGCCACAC | AGAGCAATAACCCCGTGTCC | 196 |
| *Fcgr1* | AGGTTCCTCAATGCCAAGTGA | GCGACCTCCGAATCTGAAGA | 198 |
| *Fcgr2b* | CTAGGAAGGACACTGCACCA | GACAGCAATCCCAGTGACAG | 114 |
| *Fcgr3* | TATCGGTGTCAAATGGAGCA | TATGGCACCTTAGCGTGATG | 130 |
| *Fcgr4* | TCCGGATATCTGTGGTGACA | GCTTGAATGCCAGAGAAAGC | 100 |
| *Fcgrt* | ACTGCTAGGCCACCTGGAG | AGGAGAAAGCAGCACAGGTC | 122 |
| *Fcmr* | CTTCATGAGCAAAGGACACG | AAGGTCGGAATCAGGATGTG | 101 |
| *Fcrl1* | ACCTTGGCTCCTGCTACTGA | AGCTTGTCTCCCTCCATCAC | 111 |
| *Fcrl5* | TCATTCTCACCCTGTGTGGT | ACGACCTCTCCTCGAAAGAA | 143 |
| *Fcrl6* | AGGTGGACCCCAAGAATCCAA | ACCATACACTGGTAGAGCCCA | 139 |
| *Fgfr1* | GAACGGGAGTAAGATCGGGC | TCCGATAGAGTTACCCGCCA | 160 |
| *Fgfr2* | AGTCCAGCTCCTCCATGAAC | GCGTCAGCTTATCTCTGGGG | 150 |
| *Fgfr3* | GTGCGTGTAACAGATGCTCC | CTCCGGGCGAGTCCAATAAG | 93 |
| *Fgfr4* | CCTGCCGGGATCGTGAC | CAGGATCAGGGCCAGACTTC | 87 |
| *Fgfrl1* | ACAAGGCCGGTGCCATCAAC | TGGAACGAGTCCGCTGGATT | 60 |

| | | | |
|---|---|---|---|
| *Flt1* | CGAGACAAGGGGCTCTACAC | CGATAGGACCGTCTTCCTGC | 152 |
| *Flt3* | CGCTGGGACCGCATCACA | ACAGGCAGGTCTTGGTTTGTAA | 156 |
| *Flt4* | GAGGCACTAAGACTCCAGGC | AGCTGCCTTCTGGTCTATGC | 165 |
| *Gp6* | CTTTCTTCTGTATTGGGCTGTGT | CGTCCAGCATTACTTCTTTCCA | 238 |
| *Gpa33* | CAAGGGGAGATGGTGATCGG | CTTCAGCAACAATGGCTCGC | 153 |
| *H2-Aa* | CTGACCACCATGCTCAGCCTCT | TACTGGCCAATGTCTCCAGGAG | 107 |
| *H2-Ab1* | ACCCAGCCAAGATCAAAGTGC | TGCTCCACGTGACAGGTGTAGA | 163 |
| *H2-D1* | GGAGCTATGGCCATCATTGGA | TCTCGGAGAGACATTTCAGAGC | 125 |
| *H2-Eb1* | GTCACGGTCGAGTGGA | AGTCCAGACTGTCCTTTCTG | 140 |
| *H2-K1* | CTGGAGCTGTGGTGGCTTT | GACAGATCAGAGGTCTGGGAG | 103 |
| *H2-T23* | GACTTGGAAAGCCAGGGACAT | CACGTCGCAGCCGTACATC | 121 |
| *Havcr1* | GCTGCTACTGCTCCTTGTGA | GGAAGGCAACCACGCTTAGA | 89 |
| *Havcr2* | TCCAAGAACCCTAACCACGG | CCCACTCCAATGTGGATAGCA | 150 |
| *Hepacam* | AACGCCACCCAAAATGAAGA | CGTACTGGGCTGGTGATGTTC | 135 |
| *Hepacam2* | TGGTTGGGCTCTCTCTTTGC | GGAAGCCATAGTGCACTGGA | 138 |
| *Hfe* | GTCACCGTCTGTGCCATCTT | CAGGCTGCAGATCACTCACA | 119 |
| *Icam1* | CTGTGCTTTGAGAACTGTGGC | CAGGGTGAGGTCCTTGCCTA | 129 |
| *Icam2* | ACACGGTCTTCTTTTGCCAT | CAGTGTGACTTGAGCTGGAGG | 95 |
| *Icam4* | CCTGTTATGTTGACAGTCCTCGCTTT | ACGTAGACAGCCCCCACAGC | 107 |
| *Icam5* | GATCCTAGGCATCTCAGCG | GACAGTTAGTGCTGCAGTTGAG | 113 |
| *Icos* | GGGTGTGCAGCTTTCGTTG | ATGCGTTTCCTCTGTGGGTC | 198 |
| *Icosl* | CTCCCTTGGACATCTCGTGG | TGTGGGTTTCCTGTGGGTTC | 130 |
| *Igdcc3* | GGGTGACTATGAATGTGTGGC | CCGTGTCAATGGGAACTCTGT | 209 |
| *Igdcc4* | TACACTGGACGATGTCCCC | TCCTTCCCCAAACCGTACTC | 150 |
| *Iglon5* | CAGTGTGTGAAGGTGACAACG | TACAGGATGTTGGAGCGGTTC | 91 |
| *Igsf1* | CAGAGCGAGCCTTCTACTACC | TTGAGCCAAATGCAAAGGAGC | 157 |
| *Igsf3* | CGCCAGTACCCTGGTCTCTA | GCAAGTCAAAGGAGTCGCTG | 101 |
| *Igsf5* | TCTGTTGCTGTTGTGCCTCC | CCTCATCCGAACTGTACCCG | 150 |
| *Igsf6* | CCTTCATAGTCCTCTTCAAATCAAA | TTCATCAGTGCCCTCTTGCT | 167 |
| *Igsf8* | CACAGTCTACCCCTACACGC | GGTGCTACAGATCACCGCTT | 150 |
| *Igsf9* | TGTCCGGTTCGTGCTAATCC | GATGAGGGAACCTTCTGGGC | 114 |
| *Igsf9b* | TCAGCACCCGAGGTCTTGT | GATTACGTCGCATCGCAGAAC | 107 |
| *Igsf11* | CAGGGGCGTTCTTTTACTGG | TGGCAGAAGAGCATTTAGGGG | 105 |
| *Igsf23* | ACATGAACTGTCTTCGGGAC | TTGGCTCCTTTCATTTGGGG | 130 |

| | | | |
|---|---|---|---|
| *Il1r1* | TGAAGAGCACAGAGGGGACT | GGGTCAGCTTCGATCGTCTC | 159 |
| *Il1r2* | GTGTGCCCTGACCTGAAAGA | ATCAATAGGCGTGTGGGGTC | 131 |
| *Il1rap* | GGAATTTGTGCTGCTGACGC | CCAGGAGTCGTCTGCTTTTCT | 143 |
| *Il1rapl1* | AGCTCCGATCCCACATTTGA | TTGGGCGAGCGAGTAGTTTG | 199 |
| *Il1rapl2* | ACTGGATGAAAGGCGAGAAGT | GGCGTGTTTCCGTCCATTTC | 188 |
| *Il1rl1* | CGTGTCCAACAATTGACCTG | CAAGTAGGACCTGTGTGCCC | 104 |
| *Il1rl2* | ACGCATCCAGGAAGGAATCG | GCCCTGAAGTCTGGAACTGG | 174 |
| *Il6ra* | GTCAACGCCATCTGTGAGTG | AGTCACTCTTCCCGTTGGTG | 103 |
| *Il6st* | CCAGAGCTTCGAGCCATCC | GGAAACAAGTTCGCTGCCTC | 159 |
| *Il11ra1* | CCCTTGGAGCAAGTAGCTGT | CAGGTTTTTGCGGTCCATCC | 130 |
| *Il11ra2* | CCCTTGGAGCAAGTAGCTGT | CAGGTTTTTGCGGTCCATCC | 130 |
| *Il18r1* | GCCAACGAAGAAGCCATAGAC | GAGGCGAGAACAAGCACAGTT | 110 |
| *Il18rap* | GACATCAATGTGAAGCCGGAA | GGAAGCCAAACTGTACTCTGC | 103 |
| *Ildr1* | AAGAAAGACTCGCCTCTCCG | GGGTAACGGAGGCAAACAGA | 181 |
| *Ildr2* | GTGCTGGGGGTGGATTACC | TGGGTCTCCTGACGTGTCTC | 138 |
| *Islr* | ACGTCCTGAAGGGTATCCCA | TGGGCTGGTAGCTTAGTTGC | 81 |
| *Islr2* | AACAGATGCCAGAAGCTCCC | GTGACTCCTGCTGCTCTCTG | 99 |
| *Izumo1* | GTCACTTCGGCTGAGGAT | TTTTGATGACGAAGCATCCA | 158 |
| *Jam2* | TGGCGCCCGCGTAGATG | CTTGACGGTGGTCTTTTGATGC | 126 |
| *Jam3* | CAATGCTGTCCACAAGGACG | GACAACAAGGACTCCCCCAAT | 139 |
| *Jaml* | ACTGTCATCTTGGGTGAGGAA | CACAATCAGTTTCAGGAGGCAA | 157 |
| *Kdr* | CAGGGACGGAGAAGGAGTCTG | TCTTTCTGTGTGCTGAGCTTG | 177 |
| *Kir3dl1* | TGATGGGCCCTGTGCTGATGATG | CCTCGAATTTACCACTGTGGCTCTG | 150 |
| *Kir3dl2* | TGATGGGCCCTGTGCTGATGATG | CCTCGAATTTACCACTGTGGCTCTG | 150 |
| *Kirrel* | GTGACCCTCACCTGTGTCTG | CTGTTACTCAGGACCATGTTTGA | 86 |
| *Kirrel2* | AGATCCACTTGGGCCGTAGA | GTGAACTGGAACCCTCGCTG | 183 |
| *Kirrel3* | AGAGTATTGTGTGCCGAGCC | TCCAATACCGGCTGTGGTTC | 121 |
| *Kit* | GTCCTGTTGGTCCTGCTCCG | CTGACAAAGTCGGGATCAATGC | 158 |
| *L1cam* | CACAGAGACTGAGCTGGCAA | GGTTTGTTTCCAGCGGTAGC | 81 |
| *Lag3* | AGGCCATCTCGTTCTCGTTC | TTCCTCTGAGCCGGAAATGG | 153 |
| *Lair1* | GTCTTTCCGCCCTTCTGTTCT | AGTCATCTCCAGGCCATTAGTAG | 128 |
| *Lepr* | AGTAATTGGAGCAGTCCAGCC | TTTTCGTCAGGGGCTTCCAA | 142 |
| *Lilra5* | CGGAAGGGAATCCGCACAA | CACCTCACATGAGATGGTCAC | 89 |
| *Lilrb4a* | GGACACCTTCCAAAGCCCATC | CCCTGACACCAGGTAATCACA | 83 |

| | | | |
|---|---|---|---|
| *Lilrb4b* | AGTGTCGTCACAAAAATAAGGCT | CCTGGGCGTACACAATTCCC | 122 |
| *Lingo1* | TGCCTTTCCCCTTCGACATC | AGGCAGAATAGGACAACGCC | 85 |
| *Lingo2* | CCAACACCAACCTGTCCACT | TGTAGGCGGATCAGGTCAGA | 130 |
| *Lingo3* | GCGCTGCAATCTCACATCAC | GGCCAGTTGTCGATCTCCAG | 153 |
| *Lingo4* | GGTCCCCACTCCTTTTCCTG | CCGGGAATAGTGTCCAGTCG | 127 |
| *Lrfn1* | AACTTGAACACCCTCACGCT | GACCTCAGGAACAGTCCGTC | 143 |
| *Lrfn2* | CAGAGAGGGCTGTGCTTGTG | CAGGTTGTTGACCACGAAGG | 179 |
| *Lrfn3* | GGGAGTCTTGACCTGATGCC | AGCAGGCAAAGGAGTAGTGG | 138 |
| *Lrfn4* | GCTGACCCAGTGTGGATGTT | CACAGCCCAGCAGTCTAGTG | 199 |
| *Lrfn5* | GAGCTAGACCCCAGTACGAC | TCTCAATACCCGGCTGAATGG | 140 |
| *Lrig1* | GGACGCGCAGCCTAAACC | CAACTCCTATGGAAGCAGCG | 147 |
| *Lrig2* | CCTCTCAGCAAACGCCTCA | ACTGCAATACCGCTGTCTCA | 101 |
| *Lrig3* | CAAACGCAACCCTGACTGTG | GGGCTGTCATCTTTCGTCCA | 148 |
| *Lrit1* | AACCAGCTGATGAGGCTTCC | ACCAGGTCATAGAGCCGACA | 146 |
| *Lrit2* | GGTAGGAAAGAACGTGACTCTGCA | TACCTTCTGCAATTGGCGAGGCCA | 125 |
| *Lrit3* | CGCTGCCACCAGATTTCCTA | TCGACAGCTCGATCACCTTG | 150 |
| *Lrrc4* | GAGGAGCTTGAGATGTCAGGG | CCAGTCCGTCAAAAGCATTCC | 136 |
| *Lrrc4b* | CCCTGTCAACACCCGCTAC | TCCCACCTCGATCTTTCGC | 133 |
| *Lrrc4c* | TGGCAAACACGTTGCCTATT | TGGTGTTGGTCCTTCTGGAG | 80 |
| *Lrrc24* | GGGACGCAGACACTGTTCC | CGCAGGGTATTGTTGTGCAG | 110 |
| *Lrrn1* | CTCTGCCGGTCCTCATCCTC | CACAGGTCCTTGTGGGAGTT | 144 |
| *Lrrn2* | GCGAGTGACGGACATAGGAC | CTTTGTGCACTGGACTGGGA | 93 |
| *Lrrn3* | ATGCCTCTACATCTGCAAGTCT | GATGCTTGCCGTCTCCTCAG | 150 |
| *Lsamp* | TCCTCGGCGGATGTCAAAC | TGGCTTCGTTGCTCTTCGAC | 79 |
| *Lsr* | GGGAGACTACTACCAGGGCA | TACGCCTCGTTGTTCCCATC | 141 |
| *Ly6g6f* | CTGGACGGTACTGGTGTACTG | CTCACGGTATTCCTTCCCTCTA | 199 |
| *Ly9* | GGTACAGACAAGCGGAGGTC | GAGGTGGGTGTAGGCTGATG | 136 |
| *Madcam1* | AGTACCCTACCAGCTCAGCA | TACACCCTCGTCACAGGACA | 121 |
| *Mag* | GCCTAGCAGAGAACGCCTAT | GACAGTCTCCCCCTCTACGG | 116 |
| *Mcam* | AACTGGTGTGCGTCTTCTTG | GCTTTTCCTCTCCTGGCACA | 75 |
| *Mdga1* | ACATCAGCGAGCGGGTCTA | CTGGAACTTGTCCGATGCG | 128 |
| *Mdga2* | ATCCGCTATGGTCGTCGAGT | CATCAATGGCGATGTCACCC | 138 |
| *Mertk* | GTTGGTGGATACGTGCATCTG | CTCGGTCTCTTCCCACTTCTC | 141 |
| *Mfap3* | GGAGTGTCTTCTCACGGTGG | AGAGCCCACGGTCATCAAAG | 146 |

| | | | |
|---|---|---|---|
| *Mfap3l* | CAGTTCACCCCCAGTCCAAA | GCTCCGTGGTCGTTATGTCT | 144 |
| *Mill1* | TCTGCTCCCTACCAAGATATTGA | GGAAGGGCTCATCATCGAAGT | 160 |
| *Mill2* | TGCGCTATAATGTCAGAGCCC | CTTGCACAGGTCTCATTTCCC | 169 |
| *Milr1* | GAGGATCTCCAATGCCAACGA | AGTATCCCCAGTAACAGCCC | 162 |
| *Mog* | AAGAGGCAGCAATGGAGTTGA | AAGTGCGATGAGAGTCAGCA | 77 |
| *Mpig6b* | GTCGCACAGTGCTTCAAGTG | GAGCGGAATCAGGACCTTGG | 95 |
| *Mpz* | TCTCAGGTCACGCTCTATGTC | GCCAGCAGTACCGAATCAG | 133 |
| *Mpzl1* | GAATGGGACACAAGGGAAGC | ACCTGTCCTTGTGAGTAGTGGA | 138 |
| *Mpzl2* | CTGATAAAGCCGAGGGGACA | AGCTCGAAGTGTTAGTCTGTATC | 100 |
| *Mpzl3* | CTGGCAAGAGGAACAGTAGGA | CATCGGCACTTATCTCCAAGG | 109 |
| *Mr1* | GGTGAAGCGAAGCCATACTC | TCGGCTCCTTCTGTCGAGTG | 155 |
| *Musk* | CCCTGCAAGTGAAGATGAAA | TTCAAGAACTGCGATTCTGG | 173 |
| *Mxra8* | GCTGTGGAAACTGCTGCTTCTT | AAGCTCACCACAGACTCAGACA | 105 |
| *Ncam1* | TGAGTTCAAGACACAGCCAGTCC | ATAGTGTCTGATGGGGGAGCC | 130 |
| *Ncam2* | CAAACCCACCTGCGTCAATC | TCATTGTCTGATGTGGGGGC | 130 |
| *Ncr1* | TGGTCAAAGTCGAGCAACCC | GCTTGTGGCAGTCTTCAGTTG | 138 |
| *Nectin1* | TGTGAGGCCACCAACCCTAT | TGATCCCTCCGACCACAATC | 175 |
| *Nectin2* | TCCAGATTGTCACCGACGC | CACTCGTACCCGCACATCTT | 87 |
| *Nectin3* | CACAACTTTCCTCGGCGTTTC | AACATTCTTTCCCCACACTGC | 173 |
| *Nectin4* | ATGCAGAGTCAGCACCTTCC | AGTGGTGGACCAGGATTCAG | 99 |
| *Negr1* | AGCCAGAGCCTGTCATTTCC | TGCATTCATACTCCCCAGCC | 117 |
| *Neo1* | TGCATTGCTGAGAATGATGTTGG | TGGGAGGTAATGTTGGGATGG | 91 |
| *Nfam1* | ACAGCGCTACTGCTTTGGAA | GACGCTGTAGGGACGTGTAG | 136 |
| *Nfasc* | AGTACCCAGTGCGGGAAAAG | GGGGCTTGTTGTCCTCATCA | 99 |
| *Nphs1* | GCACTTCGTGAAACCGTGAG | GGGTTAGCAGACACGGACAC | 137 |
| *Nptn* | CCCCACTTTGGCCTTTCTTG | AGAGTTGGTTTTCATTGGCCC | 125 |
| *Nrcam* | CGTGCAGAAACGGAGACTGG | TTTCACTGGAGAGCAGCACA | 155 |
| *Nrg1* | CATCCAAACCCACCACCAGA | GAGTCTGGGTGACAGTCGTG | 160 |
| *Nrg2* | TGAATGGAGGCGTGTGCTAC | AATCGCAAAGGCAGTTTCTCC | 106 |
| *Ntm* | AGGTAGACCGGAGCCTACAG | TGACTGTTCCCGAGTGATGC | 109 |
| *Ntrk1* | CGTCATGGCTGCTTTTATGG | ACTGGCGAGAAGGAGACAG | 75 |
| *Ntrk2* | CCCGGAGAACATCACGGAAA | GGTTTCTCAGCCCCACGTAA | 95 |
| *Ntrk3* | GAGTCTGATGCGAGCCCTAC | CAGCCACAGGACCCTTCATT | 181 |
| *Opcml* | CCACCCTCAGGTGTACCATAG | GTAGGCGTGTTGACCAAAATGA | 130 |

| | | | |
|---|---|---|---|
| *Oscar* | CGTGCTGACTTCACACCAAC | GGTCACGTTGATCCCAGGAG | 99 |
| *Pdcd1* | CGGTTTCAAGGCATGGTCATTGG | TCAGAGTGTCGTCCTTGCTTCC | 142 |
| *Pdcd1lg2* | GTACCGTTGCCTGGTCATCT | CCAGGACACTTCTGCTAGGG | 169 |
| *Pdgfra* | CACCGGATGGTACACTTGCT | GTCATCCCGAGAGGCACAAA | 120 |
| *Pdgfrb* | ATGGGTGGAGATTCGCAGGAG | TCGGATCTCATAGCGTGGCTTC | 156 |
| *Pecam1* | GCATCGGCAAAGTGGTCAAG | GGGTGCAGTTCCATTTTCGG | 152 |
| *Pigr* | AATCGTGACTACCACGGAGTG | ACTGAAGCAGGAATGCCGAG | 107 |
| *Pilra* | AGGCATACAGTTTTGGCAGTC | GGTGCTTGGAAGTGTAGTGGG | 85 |
| *Pilrb1* | GACCAAGCCCAGTACTTTAGTCGAG | GAGGTGACGATGAAGGGGCT | 137 |
| *Pilrb2* | GATCTTCCTGAGGTGGAAGCA | ATGCTGGTTCTCCATCCTGAC | 125 |
| *Pirb* | CAATCAGGCTGCCGAATCT | CCGCCAGAGTAGCATATACAC | 146 |
| *Procr* | GACGAAGTTTCTGCCGCTA | AGTTTTCCCAGAGAGGCGTT | 150 |
| *Prtg* | GTCCAACGAGGTGGGAGAAG | CCTGGCTTTGCTTCGGTAGA | 238 |
| *Ptgfrn* | GTGTAAGCGTGACCTGGCTA | GACACGGTAAGTCGGACACC | 145 |
| *Ptk7* | TGAACCTCGCTGCTCGTCTG | GGGAGGACGGCTCCTTGATAA | 129 |
| *Ptprd* | AAGAACACAACGACCAGCCA | CAGAAGGGGTCAGGGGTTTC | 122 |
| *Ptprf* | TGTCATGCGATCTGCCAACT | GTAGAAGGACACAGGCTCGG | 184 |
| *Ptprk* | GCCTCTTGGGATGTGGCTAAA | GCCAAATGTCGATGTAGTTGG | 151 |
| *Ptprm* | TTTCATGGGCACGCACAATC | GTGTGGGTCTCATCTGGGTA | 120 |
| *Ptprs* | AAAGCTCACCGTCCTTCGAG | TACAACCTTCAACTGGGGGC | 79 |
| *Ptprt* | CAACCGCAATGATGAAGGCT | TAGGCTGAGACAGCTCCCCA | 86 |
| *Ptpru* | GCTCAGTATGACGACTTCCAATG | TTGACCATCAAGTAGGCACCA | 95 |
| *Pvr* | CACGGTCATTGTGTGCGAAG | ACTACAGCGAGGACAAAGACG | 150 |
| *Pvrig* | CAGCAGAGGCATTGGTGTTTG | GGAAGTTGGGAAGAGTGTGGG | 172 |
| *Raet1e* | CAGGTGACCCAGGGAAGATG | CTCAACTCCTGGCACAAATCG | 79 |
| *Robo1* | GCTGGCGACATGGGATCATA | AATGTGGCGGCTCTTGAACT | 94 |
| *Robo2* | ATGATGCAGACTTGCCGAGAG | GAACTCGGACAGTGAGGGTA | 156 |
| *Robo3* | CTGGTCCCTGCTTGTTTTGTG | AGCTACCAGCGTGTATTGAC | 78 |
| *Robo4* | AATGGTGTCATCCGTGGTTAC | AGTTGGCAGCAGGCAATG | 64 |
| *Ror1* | CCTACCTCGTCCTGGAACAC | GAACCAGCGGATACTGGGAG | 153 |
| *Ror2* | AACGGGCTGAAGACCATCAC | AGTCCGGTTCCCAATGAAGC | 156 |
| *Scn1b* | GCGTCGTAGTGAGACCACC | CCCGACTACCGTTCCACAC | 158 |
| *Scn2b* | CCCTCAATCACCCTTTCCCAC | GCGCTGTGACTTCCATGCTC | 156 |
| *Scn3b* | AGATGCCTGCCTTCAACAGA | GGCTTCTGTCTCCGAGGGTA | 107 |

| | | | |
|---|---|---|---|
| *Scn4b* | TGGGAACCGAGGCAATACTC | CCAACGACAGGTACATGGGA | 83 |
| *Sdk1* | GTCGGATTTTAGGGGAAGTGGA | TAGGGTTCTGGAGTTTGCTGA | 120 |
| *Sdk2* | GCCGCTCTGGACTCTACTG | ATGGCGGTACATCATCTTGGG | 68 |
| *Sema4a* | GCCATGTGGTCATGTATCTGG | GGTTTCGAACAGGCTCAGAGT | 126 |
| *Sema4b* | AACAGCAACCTCAGCTTCTTGC | GGCCTCATCTTGGGCTAAAGTA | 240 |
| *Sema4c* | CCTCCCATCTGTATGTCTGCG | GCTGGGTCATATGGGCATTTAC | 127 |
| *Sema4d* | AAGGTGCCAGGAACAACCC | CTTCACGACGTCATGCCAAG | 79 |
| *Sema4f* | AACGGTCAGCAGCTGTAATG | AGCTCGGGAGATAATCGGCT | 89 |
| *Sema4g* | ACCGTAGCCAACAGGACAGAA | AGTTGCCACTGTGCTCCAGTT | 274 |
| *Sema7a* | CGTGTATTCGCTTGGTGACAT | GTGGGTATGGGCTGCTTTTT | 120 |
| *Sigirr* | CGTTTGGCCTGCCGTGAAG | AAGCAACTTCTCTGCCAAGGG | 123 |
| *Siglec1* | CTAGCAACACATTGGGCAAC | CCAGTACAGTGGCCTTAGCA | 63 |
| *Siglec15* | CAGCACCGAGATGTTGACGA | ACGATCGCTATGAGAGTCGC | 80 |
| *Siglece* | TGGTACAGGGAAGGAACCGA | GTGAGGGCTGTTACAACCAGA | 248 |
| *Siglecf* | CCACAGGACCACCCTCTCCTC | GGACTTTAGTTCCTGTGTCATCTCCC | 194 |
| *Siglecg* | GCTGCTACCTGATAAAGACAGTGCC | TTTCCAATTCCGAGCCAGGGACC | 226 |
| *Sirpa* | GCTGATGGCTGCTCTCTACC | GTGCAACCGTGTGGAAGATG | 76 |
| *Sirpb1a* | TGCCCTTGAGGAGAACATGG | CAAGGAGCACAGCTACAGGT | 155 |
| *Slamf1* | TCTGCGATTGCTGGCTAA | CGAGGATGCGGACACTTT | 210 |
| *Slamf6* | GGATGGTCTGGCTCTTTCCA | ACGCCATTCATTAGCTGGGG | 91 |
| *Slamf7* | AACGCTATGGCTCGTTTCTCA | TCACAGATCCATCAAGGGCAC | 118 |
| *Slamf8* | TGCTGATGGTGGATACAAGGG | GTATTGAGGGGTTGGGTCTCC | 127 |
| *Slamf9* | ACCAGTCATTCTGCCTCTATGG | CATCTCCAGAAAACCCTTTGGC | 159 |
| *Spaca6* | TTGTGGGCAGAGAGGAAACCA | AGCATCTTTGAAGGCAAGCCAA | 194 |
| *Tapbpl* | CACTAGCAGGGGAACTGTGG | GCTGTGGGAGGGTCAGTC | 136 |
| *Tarm1* | GACTTTCCCACCGCAACTGA | CACCGACCCGGATGAGATTA | 92 |
| *Tek* | TGGAGTCAGCTTGCTCCTTT | ACCTCCAGTGGATCTTGGTG | 192 |
| *Thy1* | TGCTCTCAGTCTTGCAGGTG | TGGATGGAGTTATCCTTGGTGTT | 121 |
| *Tie1* | TACATCGGAGACGCACCTTC | CCAGCACGGGGTAACTCAAG | 132 |
| *Tigit* | CTGATACAGGCTGCCTTCCT | TGGGTCACTTCAGCTGTGTC | 134 |
| *Timd4* | TGTGGGATTTGTGCTAATGGTG | CGTCTTCATCATCCCTCCCG | 158 |
| *Tmem25* | GACTGTCAATGCCTCTGACTTC | GTGGCACTTCTATCCGTGTGG | 188 |
| *Tmem81* | TTTTCAGACCCTTCCGAGCC | AGGAAGGACCCTCAGTCCAA | 146 |
| *Tmigd1* | AAGCTACAGCGCGATCAGAC | TTCCTCCACGGTTTGGAAGC | 96 |

| | | | |
|---|---|---|---|
| *Trem1* | CTGTGCGTGTTCTTTGTCTCA | CCTTCCCGTCTGGTAGTCTC | 160 |
| *Trem2* | CTTGCTGGAACCGTCACCAT | CTTGGGCACCCTCGAAACTC | 199 |
| *Treml1* | GGTCTCCCTGTTGGTTCTTCC | CCGTCTGAACTCGTGAGGAC | 154 |
| *Treml2* | TGGTGGTGGTGTTGACATTTCTTCC | ATCCAGGGTTTAGCATAGTTGCTGC | 118 |
| *Tyro3* | TCCTCCAGAACCCGTAACCA | TGCGGGCTTCACAAGAAAAC | 128 |
| *Ulbp1* | CTAACACAACCGGAAAGCCCCT | CAGTGCTTGTGTCAACACGGA | 132 |
| *Unc5a* | TGGCCTATGGGACCTTCAAC | GGCTTGTGCAGAGTGAGGTA | 130 |
| *Unc5b* | AACTGCACTGATGGGCTGTG | AGAACCGCTACCACCACAAA | 101 |
| *Unc5c* | AACAGCGAGTGGGTTCATCA | ACTCTTCGTAGTGCCTGCTG | 180 |
| *Unc5d* | AGGACTCTTTTTCTGGGCGG | TTCCAGGGGCAGATGGGATA | 92 |
| *Vcam1* | TTGCCTCGCTAGGTTACACA | AGATGGTGGTTTCCTTGGGG | 104 |
| *Vsig1* | ACCAGTGTACTGCCATCAACA | CTATCAGGGCACCGACCAAA | 111 |
| *Vsig2* | AGACCTATGGGGGTAGTGACC | AGCTCTTTGCTATGATCGGCT | 91 |
| *Vsig4* | ATGGGACTGGAAAACTTGAGGAG | CTGCAGCGGAACAAGATATAAGG | 130 |
| *Vsig8* | GCGGAGGAAAGACCAAGCC | CCATCCCCGTTGATCCTTACA | 109 |
| *Vsig10* | GTCCCAAAACCTCCAAGGCA | CCAGTTACAGGCGAGTTCCA | 169 |
| *Vsig10l* | GCTCACCCCACCAGCAAAAA | GACAGAGAGTTGGACTGGGG | 137 |
| *Vsir* | AACAACGGTTCTACGGGTCC | CGTGATGCTGTCACTGTCCT | 101 |
| *Vstm2b* | GCCCGGAGCAAGGTAACAAA | ACACCTTCGTCCTGCAATCG | 116 |
| *Vstm4* | CAACAAGTGGACAGCCTGGT | GCCAGCTTTGAGGGAGATCA | 70 |
| *Vstm5* | GGGAGAGGACCCAAGGCATC | ATGGCCGACTGTGGAATGTA | 106 |
| *Vtcn1* | ATCCGCACCTCAAAAGGCAA | CGCAGCGTAAACTCTCTGAAC | 112 |
