## Supplementary material for "IGSF3 is a homophilic cell adhesion molecule that drives lung metastasis of melanoma by promoting adhesion to vascular endothelium": Table S2

**Supplementary Table S2.** List of shRNA sequences.

| shRNA | TRC clone No. | Target sequence (5' → 3') |
|---|---|---|
| sh*Cntn1* #1 | TRCN0000039014 | GCCAAACAAATTAGACTCTAA |
| sh*Cntn1* #2 | TRCN0000039015 | GCAGCCAATCAATACCATTTA |
| sh*Hfe* #1 | TRCN0000105415 | GCTTTCTCTGTTCACTTTCTT |
| sh*Hfe* #2 | TRCN0000105416 | CGGCTTCTGGAGATATGGTTA |
| sh*Igsf11* #1 | TRCN0000250088 | GCCCGAACAGGTCATTCTTTA |
| sh*Igsf11* #2 | TRCN0000250089 | TCATCTGCCTTGCACTAATTT |
| sh*Igsf3* #1 | TRCN0000105695 | GCCTTCATAGTAACAAGGATT |
| sh*Igsf3* #2 | TRCN0000105698 | GAGTTACAGTGCCAAGATGAA |
| sh*Unc5c* #1 | TRCN0000366902 | ATCGAGCCCAACTGGCGTAAT |
| sh*Unc5c* #2 | TRCN0000376866 | CTGTCCATGGCTCGTTCTATT |
