## Supplementary material for "IGSF3 is a homophilic cell adhesion molecule that drives lung metastasis of melanoma by promoting adhesion to vascular endothelium": Table S3

**Supplementary Table S3.** List of primer sets for cloning IGSF3 and its deletion mutants.

| IGSF3 | Sense primer (5' → 3') | Antisense primer (5' → 3') |
|---|---|---|
| Full length | CACCATGAAGTGCTTTTTCCCGGT | GTCTATGGCCCCTGGATGGA |
| Extracellular domain | CACCATGAAGTGCTTTTTCCCGGT | TGCGTCGTTGGAGCAGATGA |
| Ig1–Ig2 | CACCATGAAGTGCTTTTTCCCGGT | GTCAGTTGGCTGGACGTTGA |
| Ig1–Ig4 | CACCATGAAGTGCTTTTTCCCGGT | TTCAAGAGCTGTGATGGAGA |
| Ig5–Ig8 | CACCATGAAGTGCTTTTTCCCGGTG CTGAGCTGTCTGGCTGTGCTGGGTG TGGTGTCAGCAATGGGCTTCGCAGT CACAGC | TGCGTCGTTGGAGCAGATGATG |
| Ig1–Ig5 | CACCATGAAGTGCTTTTTCCCGGT | CTGCAGCACCCTGATCTCCA |
| Ig1–Ig6 | CACCATGAAGTGCTTTTTCCCGGT | CTGTTTCACAGTGACTTCTG |
| Ig1–Ig7 | CACCATGAAGTGCTTTTTCCCGGT | TCGCATGACTGTCAGAGCTG |
